# A primer-independent DNA polymerase-based method for competent whole-genome amplification of intermediate to high GC sequences

**DOI:** 10.1101/2023.03.17.533076

**Authors:** Carlos D. Ordóñez, Carmen Mayoral-Campos, Conceição Egas, Modesto Redrejo-Rodríguez

**Affiliations:** Centro de Biología Molecular Severo Ochoa, CSIC-UAM, Madrid, Spain; Departamento de Bioquímica, Universidad Autónoma de Madrid (UAM) and Instituto de Investigaciones Biomédicas Alberto Sols (CSIC-UAM), Madrid, Spain; Center for Neuroscience and Cell Biology, University of Coimbra, Coimbra, Portugal; Biocant, Transfer Technology Association, Cantanhede, Portugal

**Keywords:** WGA, MDA, primer-independent DNA polymerase, piPolB, metagenomics, genomics

## Abstract

Multiple displacement amplification (MDA) has proven to be a useful technique for obtaining large amounts of DNA from tiny samples in genomics and metagenomics. However, MDA has limitations, such as amplification artifacts and biases that can interfere with subsequent quantitative analysis. To overcome these challenges, alternative methods and engineered DNA polymerase variants have been developed. Here, we present new MDA protocols based on the primer-independent DNA polymerase (piPolB), a replicative-like DNA polymerase endowed with DNA priming and proofreading capacities. These new methods were tested on a genomes mixture containing diverse sequences with high-GC content, followed by deep sequencing. Protocols relying on piPolB as a single enzyme cannot achieve competent amplification due to its limited processivity and the presence of *ab initio* DNA synthesis. However, an alternative method called piMDA, which combines piPolB with Φ29 DNA polymerases, allows proficient and faithful amplification of the genomes. In addition, the prior denaturation step commonly performed in MDA protocols is dispensable, resulting in a more straightforward protocol. In summary, piMDA outperforms commercial methods in the amplification of metagenomes containing high GC sequences and exhibits similar profiling, error rate, and variant determination as the non-amplified samples.

**Graphical abstract:** 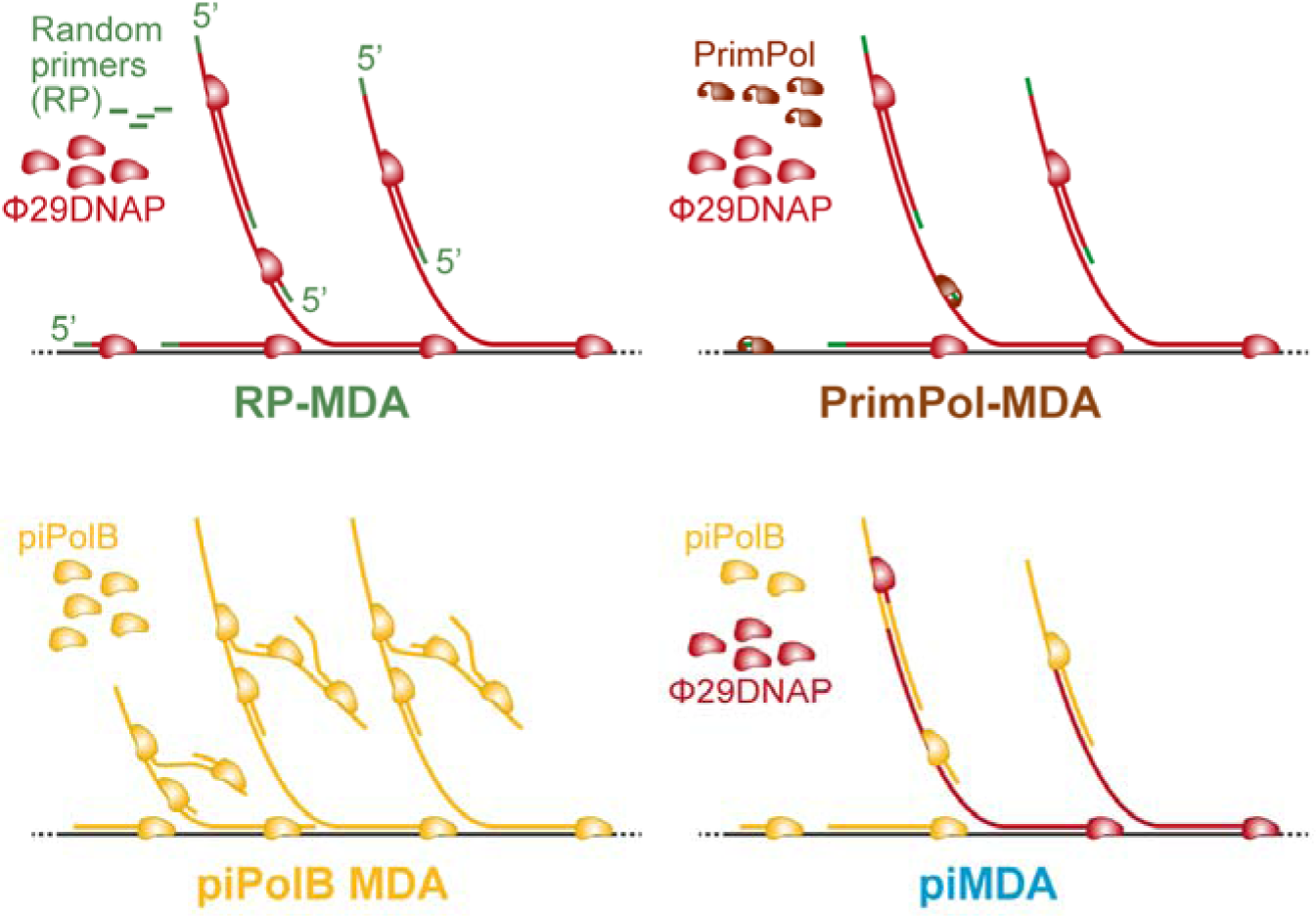

Schematic representation of methods based on multiple displacement amplification (MDA) for whole genome amplification. The diagrams above represent protocols initiated by random primers (RP-MDA) or a DNA primase-generated short DNA primers (PrimPol-MDA) and continued by Φ29DNAP, whereas the schematics below show piPolB-mediated MDA (left) and the piMDA protocol (right), in which piPolB synthesizes DNA strands that are further extended by Φ29DNAP.

## Introduction

Genomics and metagenomics have undergone a revolution in the last two decades. This is due to advances in whole-genome amplification methods and the development of massively parallel sequencing methods that have opened up previously unimagined applications in biotechnology and biomedicine (1). This has led to an increasing demand for more efficient, sensitive, and unbiased protocols aimed at developing high-throughput analysis of biodiversity and personalized medicine, among other applications.

Current technologies allow direct sequencing of DNA samples without a prior amplification step, which can reduce bias and increase data reliability. However, this is not possible when the quantity or quality of DNA is limited. Therefore, sequencing of the original raw sample is still experimental or limited to targeted testing in some scenarios (2–4). In addition, sample origin and characteristics, genome isolation procedures, and sequencing platforms can also be sources of error, bias, and underrepresentation of genomic sequences that are present in lower quantities (5–8).

To ensure sample availability for DNA sequencing, whole (meta)genome amplification (WGA), which includes PCR-based approaches, multiple displacement amplification (MDA), and a number of variations and derived methods (9–11), enables efficient amplification of minute nucleic acid samples. MDA provides faithful and reliable amplification of (meta)genomic DNA without the need for prior knowledge of the target sequence. It has been used for the amplification of complex (meta)genomic samples as well as for whole genome amplification at the single-cell level (scWGA) for almost 20 years (12–14). In addition, it is based on an isothermal protocol that simplifies its use at the point of care and promotes the development of various applications beyond genomic analysis (15–17). Most MDA protocols are based on the use of hexamer random primers (RP) and the highly processive DNA polymerase from *Bacillus virus* Φ*29* (Φ29DNAP) (15, 18, 19), which can generate very long amplicons, that allow high coverage and detection of single nucleotide polymorphisms (SNP) (20, 21). However, MDA also brings some disadvantages, such as the generation of primer-related artifacts, chimeric DNA sequences, or biased poorer amplification of sequences with extreme GC content (22–27), which may affect uniform genome coverage and sensitivity for the detection of minor alleles or underrepresented sequences (23, 28, 29). In addition, the extreme processivity of Φ29DNAP leads to the overamplification of circular DNA molecules, which compromises its use for some metagenomic applications, such as metaviromic analysis (30, 31). Accordingly, several new tools and methods have been developed to increase the accuracy of WGA. These include both engineered versions of DNA polymerases with improved MDA performance, thermostability, or tolerance to chemicals (32–34), and alternatives that combine both PCR and MDA, such as MALBAC or PicoFLEX/GenomeFlex. The latter methods have been reported to have limited coverage compared with Φ29DNAP-based MDA but offer higher single nucleotide variants (SNV) detection rates, with MALBAC having higher uniformity and a lower allelic dropout (ADO) rate in the portion of the genome covered (29, 35). However, the specialized polymerase required for this reaction lacks proofreading capacity, resulting in increased error rates (36). Another promising modification of MDA-based WGA is TruePrime, which is based on the combined use of a *Thermus thermophilus* primase-polymerase enzyme (TthPrimPol) to generate short DNA primers that are subsequently extended by Φ29DNAP (37). This protocol is expected to reduce primer-related artifacts and provide a better representation of sequences with high-GC content (38).

In this work, we have developed and assessed new MDA methods based on recombinant *Escherichia coli* primer-independent PolB (piPolB), previously characterized as having faithful DNA polymerase activity, along with proofreading, translesion synthesis, and DNA primase capacities (39). Our results have shown that the use of piPolB in the absence of DNA primers or accessory factors can lead to successful DNA amplification, although the amplified DNA product also contains a high proportion of duplicated and low complexity sequences, corresponding to spurious *ab initio* DNA synthesis (40). In contrast, the combination of piPolB with a highly processive enzyme such as Φ29DNAP, a protocol hereafter referred to as primer-independent MDA (piMDA), results in the amplification of genomic samples at a level similar to commercially available kits, which is further increased when a prior alkaline denaturation step is added (piMDA+D protocol).

Deep sequencing of MDA products from a mixture of bacterial genomes containing high-GC sequences obtained with piMDA and piMDA+D protocols shows that both alternatives achieve competent amplification and an improved assembly compared to commercial random primers and PrimPol-based MDA. Overall, our results suggest that the piPolB polymerase, primase, and proofreading capacities result in high-fidelity DNA molecules that can subsequently be extended by the highly processive Φ29DNAP for competent and accurate amplification of DNA samples. This work paves the way for the use of novel tailored MDA methods for environmental and biomedical applications.

## MATERIALS AND METHODS

### DNA amplification substrates

The single- and double-stranded M13 DNA was obtained from New England Biolabs (NEB). The genomes mixture was generated from the combination of four different bacterial genomes. *Escherichia coli JM83, Micrococcus luteus* (later corrected to *Kocuria rhizophila*) and *Pseudomonas aeruginosa* strains were taken from the teaching collection of the Molecular Biology Department at Universidad Autónoma de Madrid. *Bacillus subtilis 110NA* (41) was taken from Margarita Salas laboratory stock. Genomic DNAs were purified from 3 mL overnight cultures grew at optimum temperature (30 °C or 37 °C) by Dneasy Blood & Tissue Kits (QIAGEN). Additionally, 1 mg/mL metapolyzyme mix from a stock diluted in PBS pH 7.5 was included in the lysis step to enhance digestion. The mock metagenome sample was then prepared by mixing an equal mass of purified genomic DNAs.

Genomic reference sequences were selected from the top hits of BLASTN queries on NCBI Nucleotide Database, using a subset of the largest contigs from the control assemblies (Table S1).

### Multiple displacement amplification assays

The DNA polymerase from bacteriophage Φ29 from Margarita Salas laboratory was purified as described (42). The piPolB was purified as untagged recombinant protein as reported previously (39).

Unless otherwise stated, piPolB isothermal MDA reactions were performed employing 250 nM piPolB in the presence of 20 ng of either M13 ssDNA (m13mp18, NEB), dsDNA (m13mp18 RF I, NEB) or genomes mixture DNA. The piPolB MDA reactions contained 50 mM Tris-HCl pH 7.5, 1 mM DTT, 4% (v/v) glycerol, 0.1 mg/mL BSA, 0.05% (v/v) Tween 20, 20 mM MgCl_2_, 500 µM dNTPs, 10 mM ammonium sulfate, in a final volume of 10 μL for 30 °C for 16 h.

Characterization of piPolB amplification product was performed by digestion of the indicated volume of amplification product with diverse nucleases. Samples treated with 0.2 U/µL Micrococcal Nuclease (Worthington) were incubated at 37 °C for 2 h in 50 mM Tris-HCl pH 7.5, 1 mM DTT, 4% (v/v) glycerol, 0.1 mg/mL BSA, 0.05% (v/v) Tween 20, 25 mM CaCl_2_. 1 U/µL or the indicated concentration of EcoRI-HF, HhaI, or MnlI was at 37 °C for 2 h (EcoRI-HF) or 3 h (HhaI or MnlI) in rCutSmartTM Buffer (NEB). Nuclease S1 (Boehringer Mannheim) was tempered at 37 °C for 3 min before the incubation with the sample at 37 °C for 1 min in Reaction Buffer for S1 nuclease (Fermentas). After the digestion, enzymes were inactivated according to manufacturer’s instructions.

The so-called primer-independent DNA amplification protocol (piMDA) (see below) reactions were performed in 35 mM Tris-HCl pH 7.5, 1 mM DTT, 0.1 mg/mL BSA, 10 mM ammonium sulfate, 50 nM piPolB, 500 nM Φ29DNAP, 20 mM MgCl_2_, and 500 μM dNTPs. In the piPolB+D and piMDA+D protocols, DNA was subjected to a previous denaturation step with 75 mM NaOH in a volume of 5 μL for 3 min, and subsequently neutralized adding 5 μL of 250 mM Tris-HCl pH 7.5. Amplification reactions were carried out at 30 °C for 3 h unless other conditions are specified. Reactions were terminated by heating samples at 65 °C for 10 min. MDA controls were performed employing a random-primers-based MDA (RP-MDA) kit as random-primer-based method: Repli-g Mini Kit from QIAGEN; and a TruePrime-based-MDA (PrimPol-MDA) kit: TruePrime Whole Genome Amplification Kit from 4BaseBio. Manufacturer’s specifications were followed in commercial protocols. Furthermore, a final heating step at 65 °C for 10 min was always conducted for protein inactivation in MDA protocols. Electrophoresis analyses of amplification product were carried out by digestion of 2 μL (for amplification of circular DNA) or 5 μL (for samples from metagenomic DNA) of each reaction with 1 U/μL EcoRI or EcoRI-HF at 37 °C in a final volume of 10 μL prior to electrophoresis in 0.7% agarose 1X-TAE and visualized with ethidium bromide (Sigma-Aldrich) or GreenSafe Premium (NZYTech).

Absolute amplification measure of DNA input sample and amplification products was performed by fluorescence quantitation with the AccuBlue High Sensitivity dsDNA Quantitation Kit (Biotum). Black opaque microplates with 96 wells (Greiner) were used for accomplishing the fluorescence reactions which were measured with FLUOstar Omega (BMG Labtech).

### Amplified metagenomes sequencing and reads processing

Control genomic DNA and amplification products were ethanol-precipitated and ∼1 mg of DNA was sheared into ∼350-bp fragments using a Covaris M220 focused acoustic shearer (Covaris, Woburn, MA, USA) following the manufacturer protocol. Sequencing libraries were prepared with the TruSeq DNA PCR-Free Low Throughput Library Prep kit (Illumina) according to the manufacturer’s protocol. Samples were then sequenced in the Illumina NextSeq 550 with the 2×75bp NextSeq V2.5 High Output kit, at Genoinseq - Next Generation Sequencing Unit (Biocant, Cantanhede).

Adapters and low-quality reads were filtered with Trimmomatic v.0.35 (parameters: LEADING:3 TRAILING:3 SLIDINGWINDOW:4:20 MINLEN:35). Quality check (QC) before and after trimming was performed with Seqkit v2.3 (43) and FastQC v0.11.9 (https://www.bioinformatics.babraham.ac.uk/projects/fastqc/), and individual reports subsequently aggregated with MultiQC 1.11 (44) for comparison. Trimmomatic filtered reads were used for *de novo* profiling using Metaphlan v.4.0.0 (45) using the --unclassified_estimation option. Additionally, trimmed reads were mapped against the reference genomes with Bowtie2 v2.3.5.1 (46) and further analyzed for coverage depth, breadth, and aligning mismatches with Weesam v1.6 (https://bioinformatics.cvr.ac.uk/weesam-version-1-5/) and Alfred v0.1.16 (https://www.gear-genomics.com/alfred).

Fastp (47) was additionally used to obtain a list of overrepresented sequences in each dataset or raw reads. Identification of low-complexity sequences in Metaphlan unclassified reads was performed with BBduk (https://jgi.doe.gov/data-and-tools/software-tools/bbtools/). GC bias was assessed using Picard tool CollectGcBiasMetrics (https://broadinstitute.github.io/picard/).

Trimmomatic filtered reads from each sample were assembled with metaSPAdes (48). Genomic assemblies were analyzed as minimal metagenomes with Metaquast (49) and also annotated with Bakta (50). Comparative assessment of k-mers diversity present in the reads and in the corresponding assembly was carried out with KAT (51) using a K-mer size of 16, according to the size of the amplified metagenome, as suggested previously (52). Single-nucleotide variants (SNV) detection with Snippy 4.6.0 (https://github.com/tseemann/snippy). Finally, MetaCoAG (53) was used for *de novo* metagenomes binning (coverage >5% and p-value <10^-10^) and bins were assigned to reference genomes using Mash distances (54).

## Data analysis

Data were analyzed using R and RStudio (55) and plots were generated with ggplot2 package (56). The final figures also contain minor modifications made with Illustrator (Adobe). As indicated, analysis of inter-groups significant differences was carried out by pair-wise t-test or Wilcoxon rank tests, when normality and homogeneous variance test prescribed the use of non-parametric analyses.

The correlation between normalized coverage and frequency of sequence windows at each GC content from Picard output was analyzed using the Corrplot package (57). Outliers that would impair these analyses were removed as follows. For metagenome correlation analyses GC values below 20% and over 80% were removed. In the case of correlations for individual reference genomes (Figure S6), GC contents with a sequence windows frequency below 10 were discarded.

### Data availability

Raw sequencing data has been deposited in the e-cienciaDatos repository with DOI: 10.21950/HCNDGF

An annotated markdown report of the analysis containing the R code and plots is available in a Github repository (https://mredrejo.github.io/pimda/).

## RESULTS

### Primer-independent isothermal MDA by recombinant piPolB

The DNA priming capacity of piPolB together with a 3’-5’ exonuclease proofreading activity and strand displacement capacity (39) make it a promising candidate for use in DNA synthesis by isothermal multiple displacement amplification (MDA). As shown in Figure 1A, piPolB can amplify both single- and double-stranded DNA starting from 1 and 10 ng of input nucleic acid, respectively. Similar to other MDA protocols, piPolB MDA is affected by high ionic strength and requires magnesium as a cofactor (Figure S1A).

**Figure 1.**
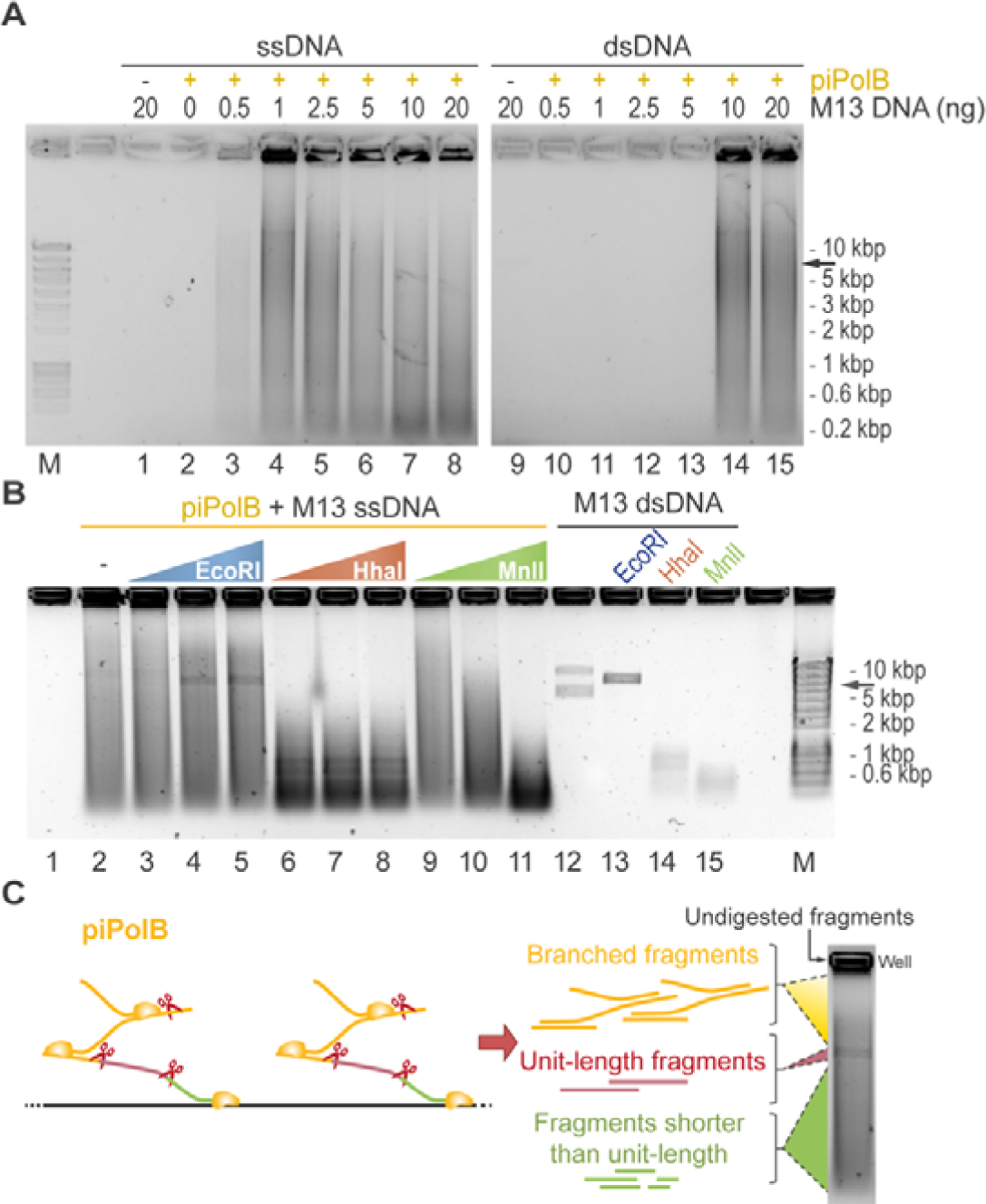
Isothermal multiple displacement amplification by piPolB (piPolB-MDA). (A) Agarose electrophoresis of EcoRI-digested product from single-stranded and double-stranded DNA amplification by piPolB. (B) Digestion analysis of amplified DNA product. Amplification reactions were performed with 1 ng input ssDNA and subsequently digested with increasing concentrations (0.07, 0.7 and 7 units) of EcoRI (single-site) and two different promiscuous restriction enzymes, HhaI (26 sites) and MnlI (62 sites). As indicated 200 ng of M13 dsDNA was also digested as a control (lanes 13-15). Lane 12 corresponds to undigested M13 dsDNA, thus containing supercoiled and relax circular forms. Migration of linear M13 is indicated with a black arrow (7.25 Kbp). (C) Schematic representation of piPolB MDA DNA product as an hyperbranched DNA structure because of multiple initiation events. Thus, single-site enzymatic digestion gives rise to a mixture of small fragments, unit-length molecules and branched molecules that move slower than unit length.

As expected, given the limited processivity of piPolB (1-6 kB, see ref. (39)), the amplified DNA product migrated as a smear in agarose electrophoresis, even after digestion with a single-site restriction enzyme that would generate unit length only if long concatemers were produced (see below). Digestion of DNA product synthetized by piPolB with different nucleases (Figure 1B and Figure S1C) indicates that piPolB synthesizes dsDNA by means of repeated initiation events that give rise to hyperbranched dsDNA structures in which fragments longer than unit length (in this case 7.25 kb) would be a minority (Figure 1C).

To obtain larger DNA products we then combined piPolB with Φ29DNAP (Figure 2). As expected, Φ29DNAP can extend DNA fragments generated by piPolB, resulting in long amplicons that in the case of plasmid substrates are made up of concatemers that can be digested into unit-length fragments. This new protocol also allows the amplification of smaller amounts of DNA, especially for dsDNA (Figure 2B). Amplification of dsDNA (plasmid or metagenome) is impaired with increasing piPolB concentration, likely due to competition between the two DNA polymerases for the free 3’OH ends, which would be more scarce for dsDNA than for ssDNA. However, the amplification yield correlates with Φ29DNAP concentration, suggesting that it is responsible for the main amplification product. This combined protocol of piPolB+Φ29DNAP in a single step will be referred to hereafter as piMDA or piMDA+D, when a denaturation step was performed previously.

**Figure 2.**
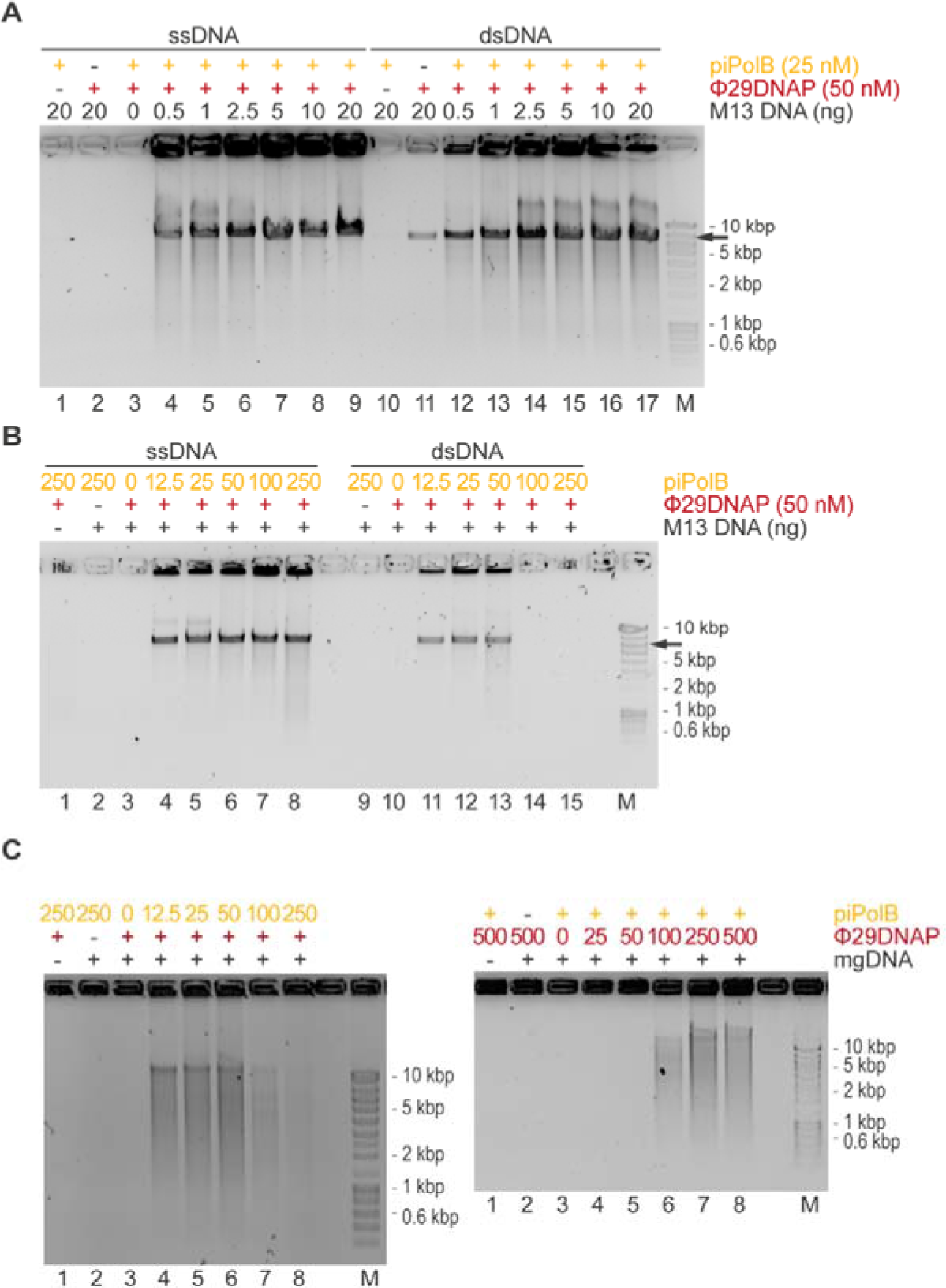
Proficient isothermal DNA amplification by piPolB coupled to Φ29DNAP (piMDA) generates long amplicons. (A) EcoRI-digested DNA products of piMDA product on different DNA substrates reveals the formation of long dsDNA amplicons, larger than the M13 genome unit, increasing the sensitivity for amplification of ssDNA (lanes 1-9) and, especially, dsDNA substrate (lanes 10-17). The ratio of piPolB: Φ29DNAP affects the DNA yield on DNA amplification of plasmid (M13, black arrow). (B) and a mock metagenomic DNA (mgDNA, C).

To evaluate the amplification efficiency of the piPolB MDA and piMDA methods in detail, we prepared a genomic DNA sample (see Methods) consisting of four genomes from Gram-positive and Gram-negative bacteria, of different sizes (2.7-6.5 MB) and moderate to high (45-70%) GC content. The final sample (Table S1) is approximately 18 MB long and has an average GC content of 58%. This sample was used for comparative MDA assays with piPolB-based protocols and two commercially available kits based on Φ29DNAP, namely RepliG (Qiagen) for Random-Primers MDA (RP-MDA) and TruePrime (4BaseBio) for a primase-based MDA (PrimPol-MDA). The results (Figure 3 and Figure S2) show that the piPolB MDA protocol is comparable to the available MDA alternatives in terms of DNA yield, but the yield of the piMDA method, especially the version with a DNA denaturation step (piMDA+D), outperforms the classical, random-primers based MDA (RP-MDA) in amplifying of our genomic sample. Under these conditions, amplification using the RP-MDA method was significantly less productive than the other methods, likely due to the abundance of high-GC sequences in the analyzed sample.

**Figure 3.**
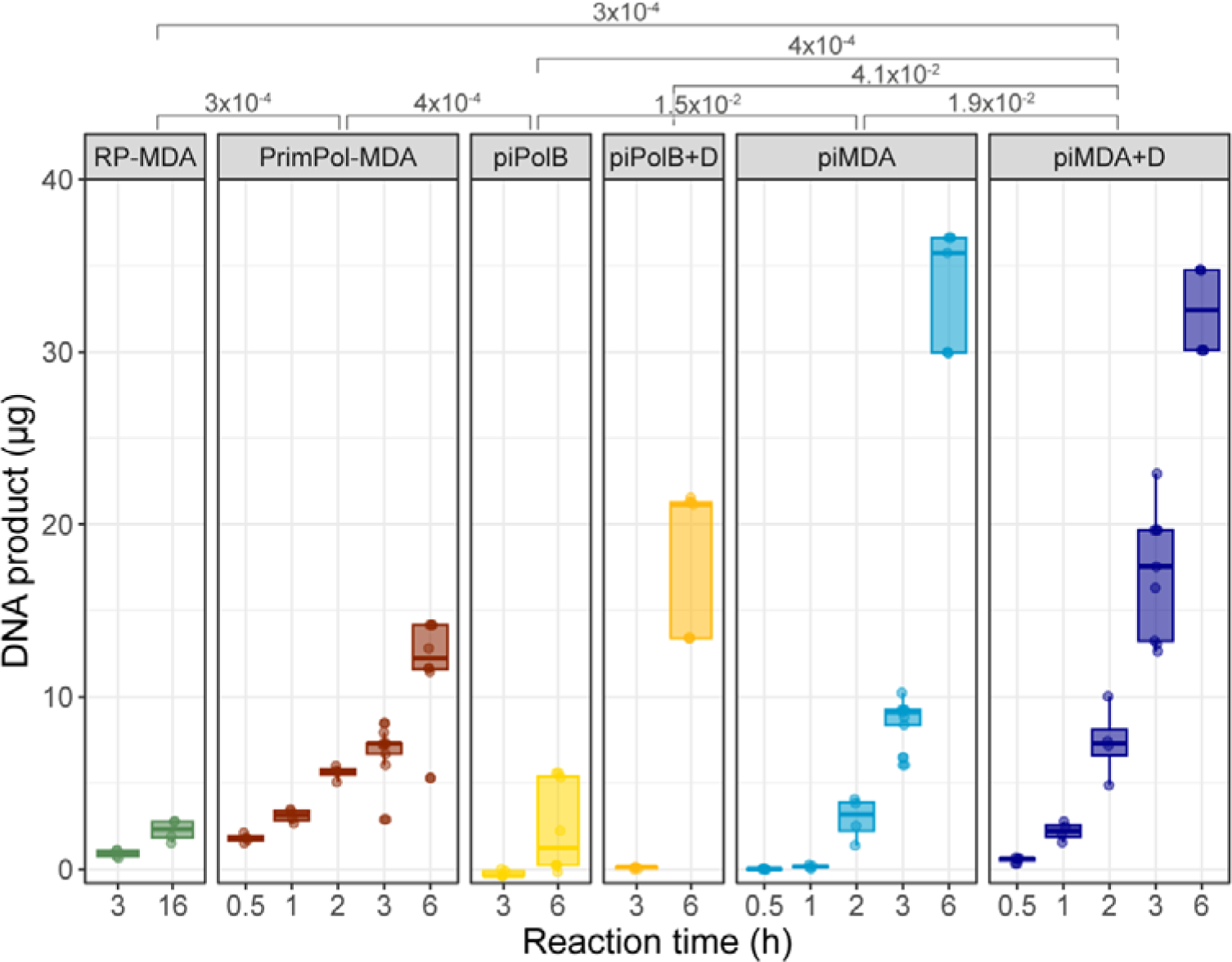
Time-course assays of different DNA amplification protocols. Quantification of piPolB-based MDA protocols and commercially available random-primers MDA and PrimPol-MDA DNA products. Amplification assays with commercial kits were performed following manufacturer’s instructions. When indicated (+D), a prior alkaline denaturation step was performed for piPolB and piMDA protocols. Four independent MDA experiments with 0.4 ng/µl genomic DNA input were quantified after the indicated amplification time at 30 °C. Significant p-values from unpaired Wilcoxon rank sum tests for pairwise comparisons of DNA product/hour are indicated above the graph.

### High sequencing depth underscores the need for a highly processive DNA polymerase in MDA

Samples from MDA experiments of the genomes mixture were sequenced using a PCR-free library preparation and short reads paired-end strategy to minimize the sequencing bias (58). The amplification products of piMDA, piMDA+D, and RP-MDA and the nonamplified control samples (NA) were analyzed in duplicates from independent amplification experiments (Table S2). Deep sequencing yielded an average total length of more than 280X of the input metagenome and provided high sequence coverage for all samples.

Adaptor trimming and quality filtering revealed the first differences between samples, with reads from piPolB and piPolB+D MDA products having lower average quality scores (Table S2) and a higher proportion of reads discarded at this stage (Table S3). This suggests that library generation was defective in these samples, likely due to the hyperbranched structure of the amplified DNA product generated by piPolB. This could hinder the generation of a homogeneous DNA shearing and adapter ligation, making detailed evaluation of the piPolB and piPolB+D MDA methods difficult.

As expected for DNA sequences obtained by MDA, the quality assessment also showed that all samples contained more over-duplicated reads (duplication level > 3-4) than the non-amplified control samples (Figure 4A). Importantly, the proportion of duplicated reads was lower for the piMDA and piMDA+D samples than for RP-MDA and PrimPol-MDA up to a duplication level ≤10, suggesting that the latter protocols are more prone to overduplication of some sequences. However, the piPolB and piPolB+D samples contained a higher proportion of highly duplicated sequences. This pattern would be consistent with repetitive initiation events followed by distributive strand elongation, resulting in a biased amplification product.

**Figure 4.**
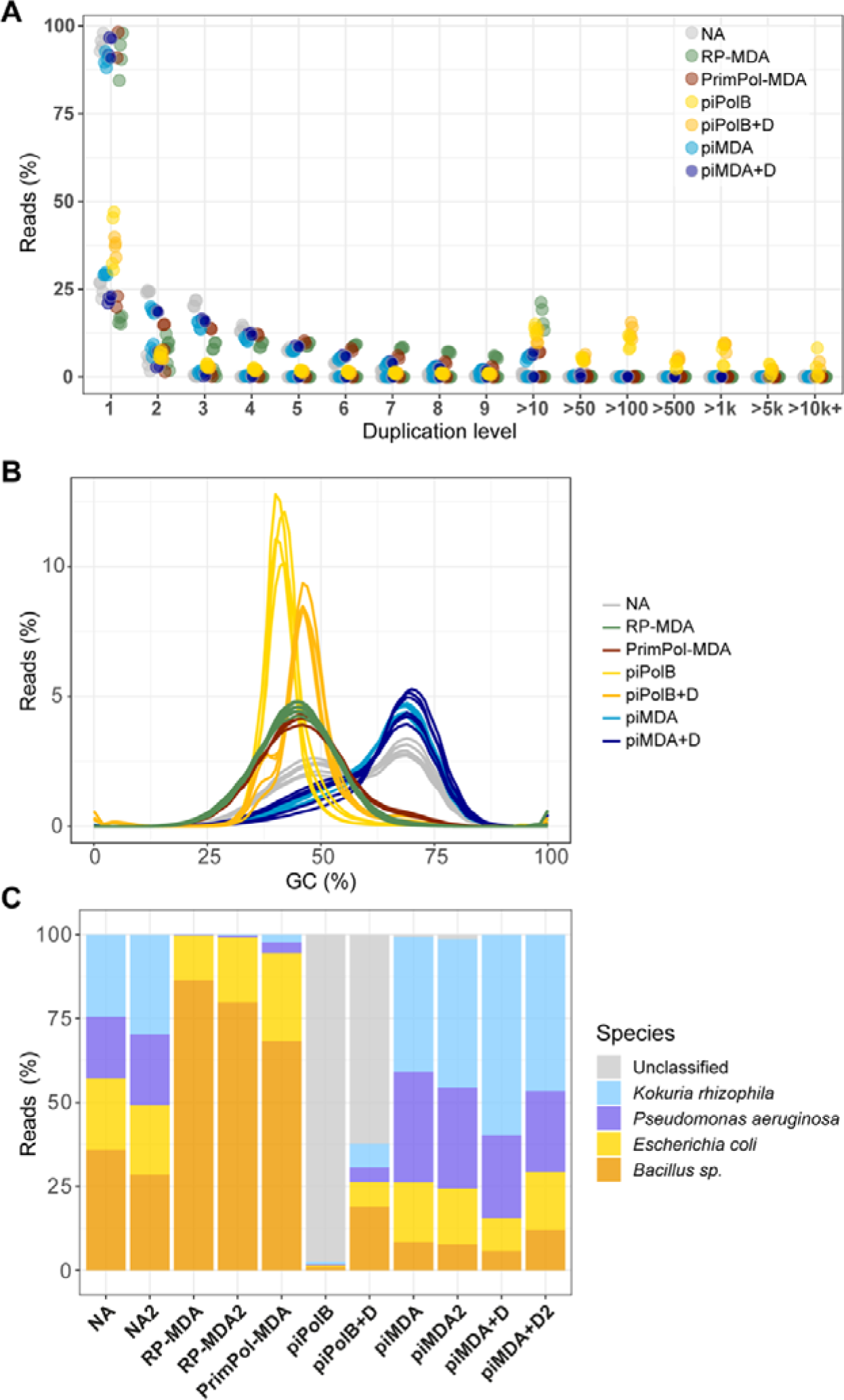
Characterization of the reads from each MDA experiment. Trimmomatic output files containing paired and unpaired reads from forward and reverse raw reads were analyzed with FastQC and subsequently evaluated for metagenome profiling with Metaphlan4. (A) The relative level of duplication found in each sequence. The piPolB and piPolB+D overrepresented sequences were also extracted with Fastp (see Tables S9-S12). (B) GC content proportion per sample. (C) Profiling of reads across the Metaphlan database at the species or genus level. Unclassified reads are also indicated. For simplicity, samples in A and B are colored by MDA method, although paired and unpaired reads for each of the eleven samples sequenced are represented.

### Sequence coverage bias is highly dependent on GC-content

The GC content of the input genomic mixture is 57.70% (Table S1), which is very similar to the value obtained in the non-amplified samples, rendering curves with two peaks, one around 45% GC, which would correspond to the genomes of *E. coli* and *B. subtilis* and the second up to ∼68% GC, corresponding to the GC content of *P. aeruginosa* and *K. rhizophila* (Figure 4B). However, the average GC content in RP-MDA (∼44%), PrimPol-MDA (∼46%), piPolB+D MDA(∼46%), and piPolB MDA (∼42%) is reduced because the peak of high-GC content is absent from the curves, indicating a strong negative GC bias. On the other hand, the reads from the piMDA and piMDA+D products show a positive GC bias as their average GC content ranges from 60 to 65% and shows curves with a main peak at 70% GC and a shoulder around 50% GC.

We then performed the profiling of the raw data of each sample against a recently curated microbial database using Metaphlan (45). As shown in Figure 4C, reads from the nonamplified control samples show an overall even distribution among the four species that make up the mixture, with small differences between duplicates resulting in a slight underrepresentation of the larger genome of *P. aeruginosa* and an overrepresentation of the moderate-GC sequences of the *B. subtilis* genome, consistent with the expected bias and variability resulting from library preparation and short reads sequencing (58, 59). However, the random-primers- and PrimPol-based MDA promoted a very strong underrepresentation of high-GC genomes (*K. rhizophila* and *P. aeruginosa*) and, in turn, an overrepresentation of *E. coli* and especially *Bacillus* sequences. In contrast, the DNA samples generated with the piMDA and piMDA+D protocols contain some overrepresentation of genomes with high-GC content compared to the NA samples, although they are distributed across the four species and show the highest relatedness to the non-amplified samples (Figure S3). Strikingly, the samples from piPolB and piPolB+D MDA contain a high proportion of unknown DNA sequences. The proportion of unknown DNA sequences decreases with the addition of the previous DNA denaturation step in piPolB+D MDA, suggesting that amplification specificity increases when access to the DNA priming substrate is favored by double helix denaturation. Accordingly, a negligible number of unclassified reads in piMDA reads (around 1%) is absent in the samples obtained piMDA+D protocol. Importantly, the fact that these unknown sequences have no similarities in the metaphlan database rules out the presence of contaminated DNA in any of the samples analyzed.

The poor quality control and profiling results of the sequencing data of piPolB and piPolB+D DNA amplification suggest that piPolB may be able to perform *ab initio* DNA synthesis, independent of primers and template. *Ab initio* DNA synthesis has been described in various DNA polymerases since the 1960-1970s, and its contribution *in vivo* and *in vitro* to the bulk of DNA synthesis is controversial (40). The mechanism of generation of these undirected DNA products is poorly understood, but the characterization of the *ab initio* activity of Bst DNA polymerase indicated that it is strongly stimulated by DNA nickases and it results in low-GC content DNA enriched in non-palindromic repetitive sequences (40, 60). In the case of piPolB, prolonged incubations of MDA assays in the absence of input DNA (Figure S4), confirmed the spurious amplification. Extraction of the overrepresented in forward and reverse reads of piPolB and piPolB+D amplified samples shows that they consist of similar repetitive sequences (Tables S9-S12). Furthermore, examination of the unmapped reads from the Metaphlan database shows that the addition of a denaturation step not only increases mapping efficiency but also decreases the proportion of low complexity reads, from 12.6% in piPolB MDA to 0.25% in piPolB+D MDA, similar to the level in any of the other samples (Table S4). These results, together with a lower GC content in piPolB than in piPolB+D raw reads (Figure 4B) but not in the mapped reads (see below and Figure 6E), points to *ab initio* DNA synthesis by piPolB that would be reduced when access to single-stranded DNA is facilitated by the denaturation step.

In conclusion, the use of piPolB for MDA as a single enzyme is hampered by the limited processivity of the enzyme as well as by spurious DNA synthesis, whereas its combination with Φ29DNAP reduces or eliminates *ab initio* DNA synthesis and allows proficient whole genome amplification with a sequence profiling similar to that of the unamplified sample.

### GC bias affects coverage depth, breadth, and amplification fidelity

We then analyzed the mapping, coverage, and alignment mismatch statistics of each sample with the independent reference genomes in our genomic mixture. As a reference, Figure 5 shows the coverage plots for the *E. coli* (A) and *K. rhizophila* (B) genomes. The control NA samples show the most consistent coverage, although they contain spikes of overrepresented sequences that range up to 10-fold the average depth. Mapping of reads from the RP-MDA and PrimPol-MDA samples shows a jittered but overall uniform plot in the *E. coli* genome, but coverage of *K. rhizophila* is scattered with some regions of great depth coverage. In the case of the piPolB and piPolB+D samples, the plots indicate an uneven coverage depth, with peaks of strongly overrepresented sequences and a low average depth, resulting in a high standard deviation of depth, especially in the piPolB reads (Figure 6C). As for the piMDA and piMDA+D reads, the mapped reads show a high jitter but an overall similar profile for all reference genomes with few large peaks of overrepresented sequences but with high coverage of reference genomes with high-GC.

**Figure 5.**
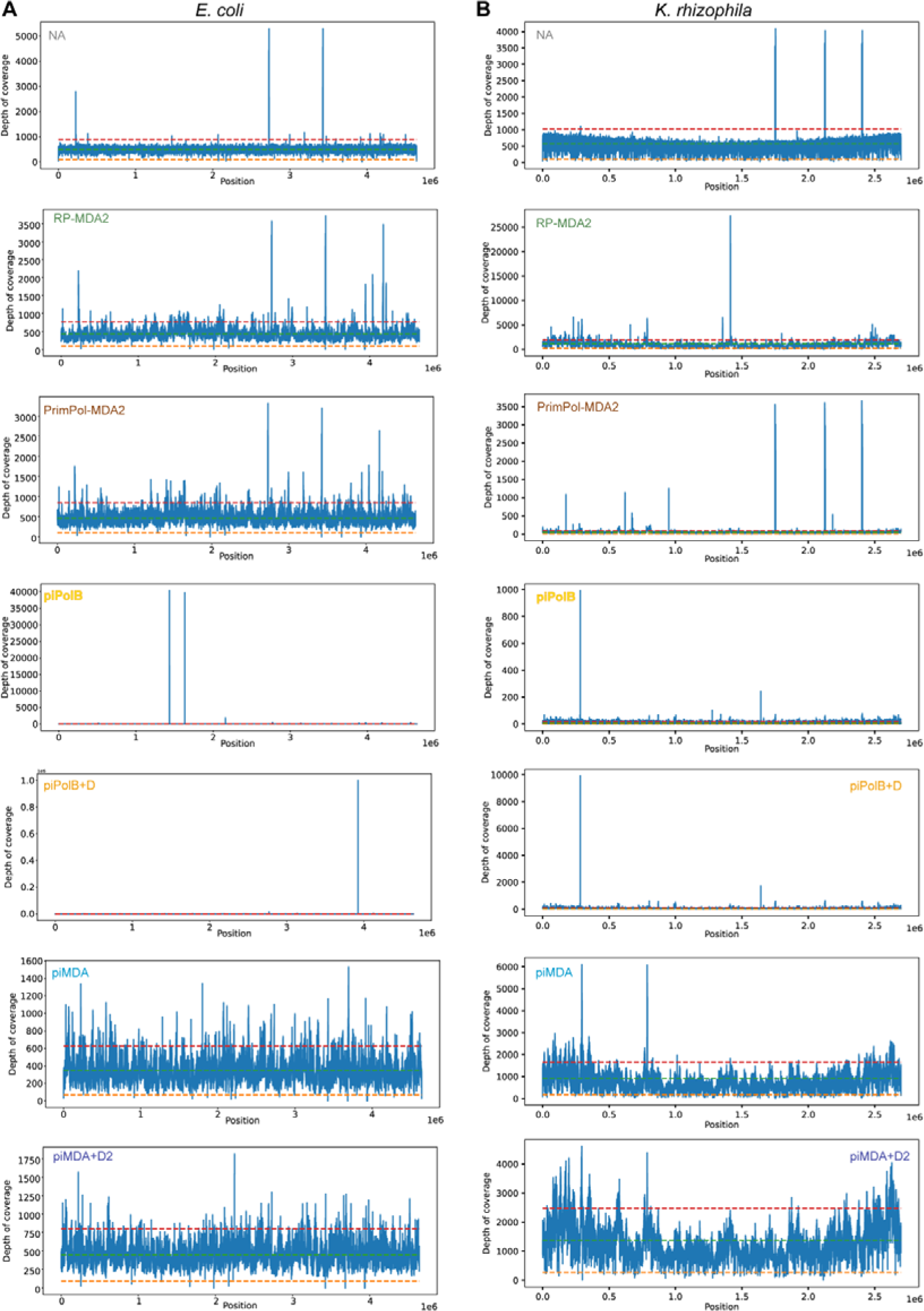
Mapping coverage of sequenced samples across the reference genomes of *E coli* (A, 50% GC) and *K. rhizophila* (B, 70% GC). Average coverage depth, x0.2, and x1.8 ranks are indicated with a green, orange, and red dotted line, respectively. Plots were generated using weeSAM (see Methods). Note that a different Y-axis scale was calculated for each sample, see Figure 8 for a coverage plot with a constant scale per sample. As indicated, mapping coverage is shown for one sample of each protocol. A full report with all samples mapping over the four reference genomes is available at the online Github repository (https://mredrejo.github.io/pimda/weeSam/weesam_full.html).

**Figure 6.**
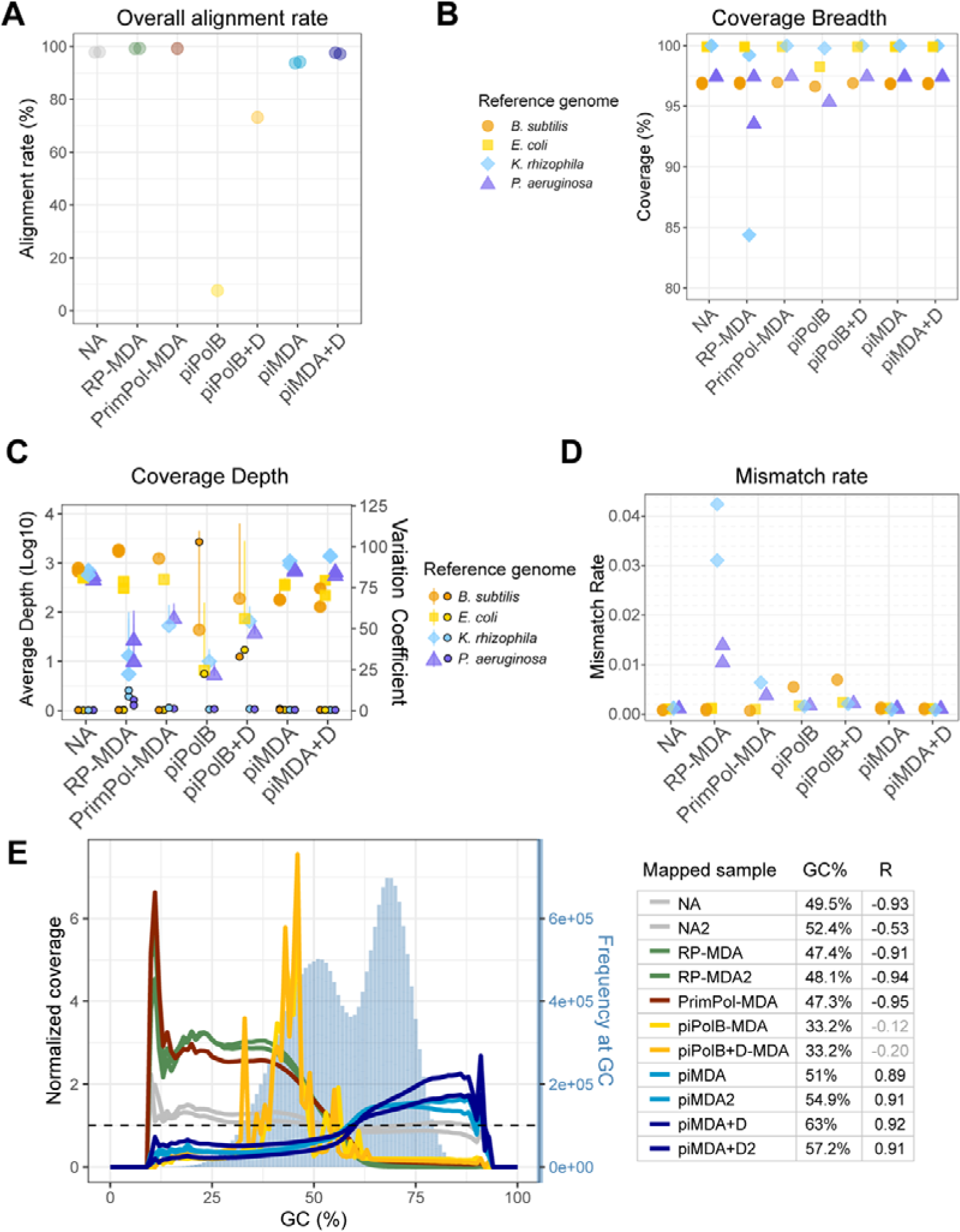
Coverage of reference genomes mixture by different MDA protocols. Mapping reads rate (A), coverage breadth (B), and average depth ± standard deviation (C) were analyzed using Weesam v1.6. The small points in panel C represent the coverage variation coefficient per reference genome (right axis). The dashed horizontal line marks the value 1 for a homogeneous coefficient of variation. The overall mismatch rate (D) was obtained using Alfred v.0.1.16. (E) Normalized sequence coverage by sample reads at a given GC content was determined using Picard v2.25.0. Percentages of GC in the reference sequence are determined from sequence bins of 100 bases, whose abundance is shown as steel blue bars (right axis). The horizontal line represents the reference value of homogeneous coverage at all GC content values. The average GC content of each sample and the correlation of normalized coverages to sequence abundance at each GC content (95% confidence interval) are also shown in the right panel. Nonsignificant R coefficients (p>0.05) are shown in grey.

The proportion of mapped reads (Figure 6A) agrees with the profiling results (Figure 4C). We also analyzed the Weesam coefficient of coverage variation, which indicates the variability of coverage depth and breadth, with values > 1 (Figure 6C) considered indicative of non-uniform coverage. High-GC genomes of *P. aeruginosa* and *K. rhizophila* are covered by RP-MDA, PrimPol-MDA, piPolB-MDA, and piPolB+D-MDA with lower depth, with the reads from both RP-MDA samples with a distinct higher coefficient of variation (Figure 6C). This is consistent with low depth and breadth (Figure 5 and Figure 6B) of the high-GC reference genomes and also with previous reports (61). This less proficient amplification of high-GC sequences is also reflected in a higher mismatch rate of the RP-MDA and PrimPol-MDA methods for these genomes (Figure 6D).

However, the samples generated by the piPolB and piPolB+D MDA protocols include a lower amplification fidelity of *B. subtilis* reference sequences that also exhibits higher depth standard deviation suggesting that overamplification may be the reason for the lower replication fidelity. In contrast, the piMDA(+D) samples show a more uniform average coverage depth of the four genomes, although with a lower depth for the *E. coli* and *B. subtilis* reference genomes and a similar mismatch rate for all reference genomes as the control samples. Detailed analysis of GC bias in the genomes mixture (Figure 6E and Figure S5) highlights the differences in high-GC content samples, with a significant negative correlation of coverage per GC content not only in the RP-MDA and PrimPol-MDA samples but also in the NA control samples, indicating a negative effect of the sequencing method in our sample, likely at the library preparation step (58, 62). Thus, the normalized coverage in the control NA samples decreases from ∼1.2 to ∼0.8-0.9 at GC content above 50-55%. In line with previous results, the piPolB(+D) amplification gives rise to scattered peaks of highly overrepresented sequences encompassing 30-50% GC content.

Previous analysis of GC bias induced by PrimPol-MDA method shows some discrepancies that might be due to the different nature of the amplified sample, as it was reported to have a minor negative GC bias, similar to that of RP-MDA in human DNA (37) but it outperformed the classic MDA protocol on metavirome amplification (38). In our genomes, we found that PrimPol-MDA exhibits strong negative GC bias, only slightly less pronounced than the RP-MDA samples.

Thus, both sets of samples show a statistically similar (Table S5) normalized coverage of over 2.5 at low-GC content, dropping to 0.1-0.2 at a high-GC content of >60%. Contrary to that and consistent with the profiling results, the piMDA and piMDA+D assays promoted a significant bias toward high-GC sequences, but with a more balanced normalized coverage, ranging from 0.4-0.5 at 45% GC content (*B. subtilis*) to 1.4-1.7 at 70% GC (*K. rhizophila*). The analysis of GC bias per reference genome suggests that bias is higher in intermediate-GC genomes, and it also reveals certain differences between duplicates (Figures S5 & S6). Nevertheless, pairwise comparisons of the normalized coverage of all samples show that piMDA+D samples are not statistically different from the non-amplified samples (Table S5).

All in all, the piMDA and piMDA+D methods provide reasonable coverage at any GC value, with higher coverage for sequences with high-GC content.

### Extraction of high-quality genomes from high-GC bacterial genomes amplified with piMDA

We then attempted to assemble the reads to compare the assembled sequences and considered them as a minimal metagenome and evaluate the extraction of individual genomes. We could obtain *de novo* assemblies with Metaspades, except for reads from the piPolB MDA sample, which could not be assembled due to the abundance of low-complexity sequences. The assembly statistics (Figure 7 and Table S6) and circular visualization (Figure 8) show that the obtained assemblies appear to be different from the reference genome for all samples of the *P. aeruginosa* genome. On the other hand, the obtained *B. subtilis* 110NA assemblies in all samples have a gap corresponding to the expected deletion across the Spo0A gene (41). In contrast, the assemblies of *E. coli* and *K. rhizophila* match the reference genomes very well in most samples.

**Figure 7.**
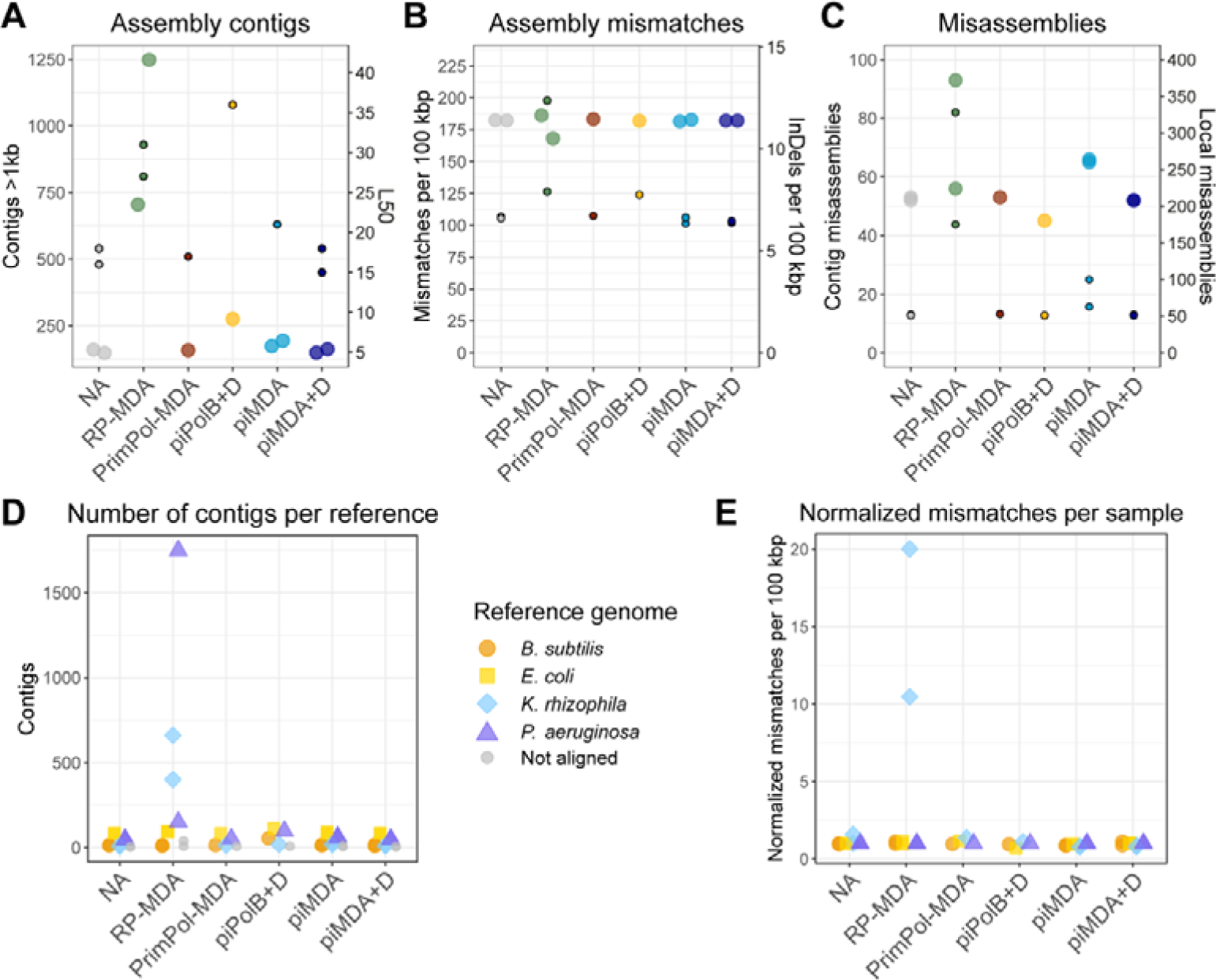
Assembly of amplified sequences. Metaspades (v. 3.13.1) *de novo-generated*assemblies were analyzed with MetaQuast v. 5.1.0rc1 using the indicated genomes as references (Table S1). (A) Number of contigs (large points) and the L50 (small points), i.e., the smallest number of contigs whose length sum makes up half of the genome size. (B) Mismatches (big points) and the number of insertions and deletions (small points) per 100 kbp. (C) Total number of misassemblies (big points) and local misassemblies (small points). (D) Number of contigs per reference genome. Non-aligned contigs are also shown. (E) Mismatches per genome and per 100 kbp are represented normalized to the control NA sample.

**Figure 8.**
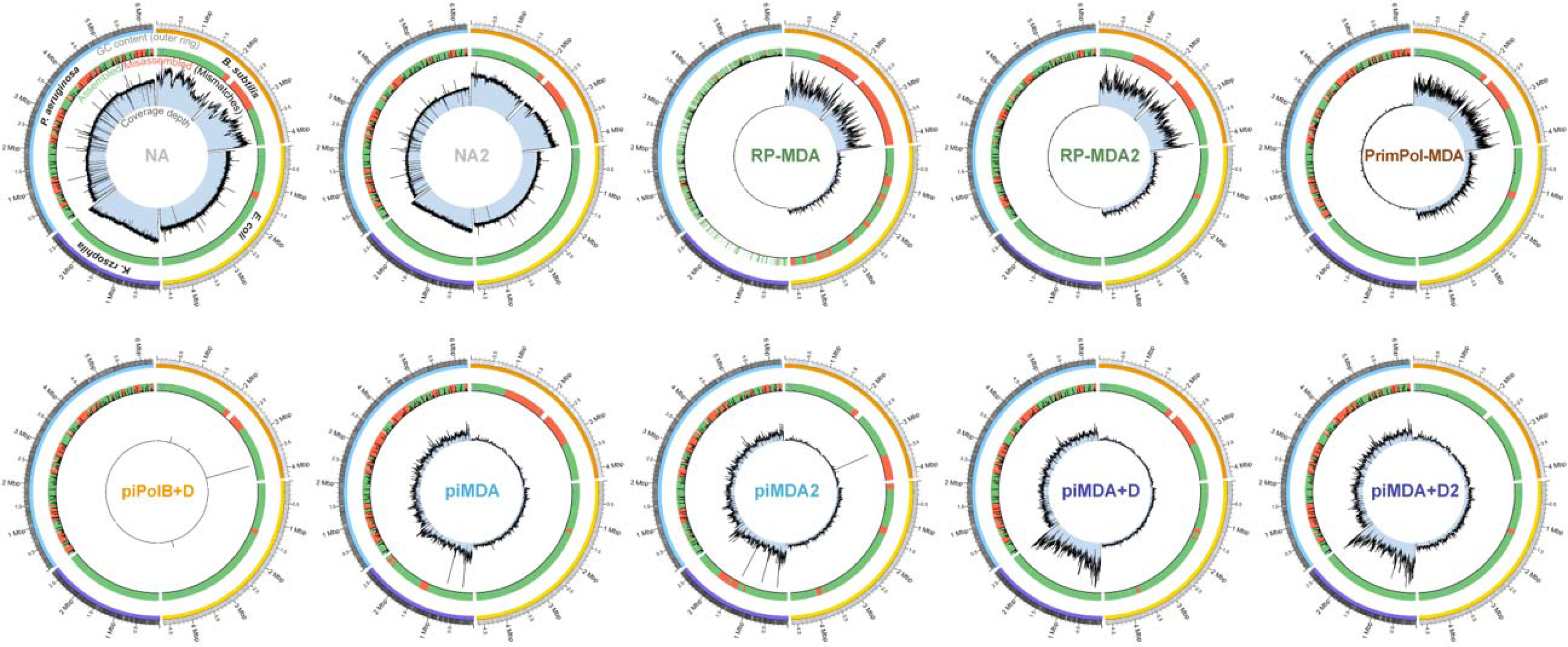
Circos plot of assemblies mapped against the reference genomes. As indicated on the NA control sample, the outer circle represents reference sequences with GC (%) heatmap [from 25% (white) to 79% (black)]. Yellow, orange, cyan, and purple color bars help to distinguish between the reference genomes of *B. subtilis, E. coli, P. aeruginosa, and K. rhizophila*, respectively. Green/red bars in the second circle represent correctly assembled and misassembled contigs, respectively, and black bars indicate mismatches. The inner circle contains a plot of the reads coverage depth, with a uniform scale for each sample in the different references.

The number and length of contigs in the final assemblies were similar in NA, PrimPol-MDA, piMDA, and piMDA+D samples (Table S6 and Figure 7). The assembly of piPolB+D sample, and its reads were also assembled in a higher number of contigs and the L50 was the largest obtained, which also indicates the presence of several small contigs, mostly from *E. coli* and *B. subtilis*. The RP-MDA assembly has a very high number of contigs (Figure 7A), suggesting many small contigs with high-GC content (> 50%, see Figure S7) mapped to the genomes of *P. aeruginosa* and *K. rhizophila*. Accordingly, this assembly also had a higher number of misassemblies and mismatches, mainly related to the *K. rhizophila* sequences. The assembly of the PrimPol-MDA sample also contains a large number of small contigs with very high-GC content (<60%), but the assembly show a better quality. K-mer comparison of reads and assemblies indicated that, as expected, only the piPolB+D sample contain a high number of k-mers excluded from the assembly and RP-MDA, RP-MDA2, PrimPol-MDA and piPolB+D samples contain k-mers in the reads occurring >2 times in the assembly, in agreement with a higher level of biased overamplification (Figure S8). Finally, the annotation of all the sequences was very similar, except for those from RP-MDA and piPolB+D amplified DNA, which span lower coding density and a higher number of pseudogenes, most likely as a consequence of poor assembly (Table S7).

For short variant calling, we used the Snippy caller, which was recently benchmarked as the best variant caller for microbial genomes and showed both strong and consistent performance across species (63). The capacity of variant calling varied widely among samples (Table S8). For example, RP-MDA and piPolB(+D) amplification result in poor variant detection of both single nucleotide polymorphisms (SNPs) and complex variants. It should be noted that with the exception of samples NA and NA2, which can call the same SNVs, there are differences between the duplicates and the second batch of sample sequencing in the case of RP-MDA, piMDA and especially piMDA+D allow better variant detection, with piMDA+D2 being identical to the control samples. Nevertheless, the PrimPol-MDA and piMDA(+D) methods show similar and good performance in variant detection, with up to 60-80% of variants coinciding with the non-amplified samples.

We then performed reference-free contig binning of the assemblies to test whether amplification affected the extraction of the metagenome-assembled genomes (Figure 9). We used MetaCoAG because it generates contig bins with abundance, base composition, and connectivity information from assembly graphs to produce high-quality bins (53), which we then assigned to the corresponding reference genome. As expected from the high coverage breadth and the statistics of the assemblies, the *E. coli* genome in all the samples can be assigned to a single bin covering 95-98% of the reference genome, whereas the genomes of *B. subtilis* and *P. aeruginosa* were more unevenly covered by the obtained bins. Consistent with the above results, the contigs of the RP-MDA assemblies result only in bins covering half of the genomes of *P. aeruginosa* and *K. rhizophila*. Interestingly, all other assemblies yield a similar binning profile, even the piPolB+D assembly, although supported by an overall lower coverage depth and breadth.

**Figure 9.**
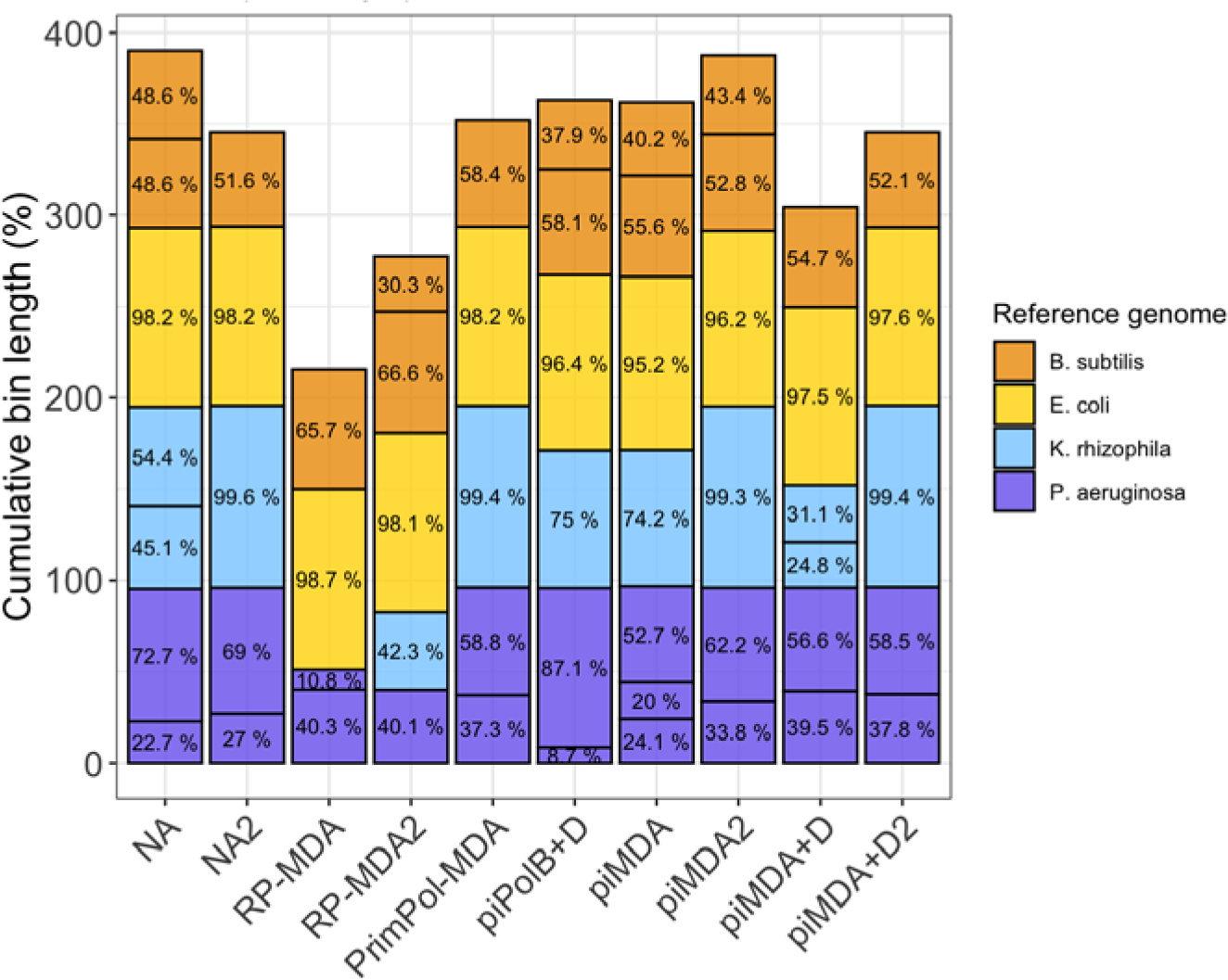
Sample binning for each amplification protocol. Bins were reference-free extracted with MetaCoAG (–mg_threshold 0.4 –bin_mg_threshold 0.2) and subsequently assigned to a reference genome using the smallest Mash distance. The proportion (%) of each bin spanned in its reference genome is indicated and it is represented accumulated in the Y axis up to the full genomes mixture (up to 400%).

## DISCUSSION

### Performance of piPolB in alternative MDA methods

The combination of 5’-3’ DNA polymerase and DNA primase activities, coupled with 3’-5’ proofreading, together with the ability for translesion synthesis and strand displacement capacity, suggests that the recombinant *E. coli* piPolB has potential for its application in novel MDA methods (39). Here we developed new MDA protocols consisting of the use of piPolB as a single enzyme or coupled with another faithful, highly processive DNA polymerase, such as Φ29DNAP. The performance of these new methods was analyzed in comparison with commercially available MDA kits for amplification of genomic sequences with moderate (45%) to high-GC content (70%), to evaluate the GC bias of each method, which has been described in detail as a major source of bias in MDA as well as in short-read high-throughput sequencing (26, 28, 38, 59, 62). We performed a detailed comparative analysis of the short reads sequenced samples, either unamplified or after isothermal 30 °C amplification with different piPolB-based protocols. Sequencing of DNA products synthesized with piPolB MDA has revealed two major drawbacks to using this DNA polymerase as a single synthesizing enzyme. First, the piPolB DNA product appears to have a hyperbranched structure resulting from repeated priming events, which may impair competent WGA and also hinders sequencing library preparation. We also detected a high proportion of unknown DNA sequences. Random primers and the presence of preexisting contaminant DNA have been referred to as the main sources of artifacts in MDA (24, 25, 27, 64), but that spurious DNA has not yet been analyzed in detail by high-throughput sequencing. In the case of the piPolB MDA products, there are no RPs in the reaction and high-throughput sequencing ruled out contamination for the not assigned reads during Metaphlan profiling. Therefore, we hypothesized that piPolB is capable of synthesizing DNA in a template-independent manner, commonly referred to as *ab initio* DNA synthesis. Although neglected or controversial in the literature, *ab initio* DNA synthesis has been reported for several DNA polymerases, particularly those of ancient origin (65–68), as is likely the case for piPolB (39). Consistent with the literature, these spurious DNA products generated by piPolB consist of low-complexity, repetitive, and low-GC DNA sequences (Tables S9-S12). Overall, these results highlight the limitations of piPolB for MDA, but also caution against the presence of *ab initio* DNA synthesis in whole-genome amplification reactions under certain reactions, that may be overlooked in MDA products. We believe that this should be further addressed in the future by developing modified protocols that could control this activity using SSBs or other factors that can prevent the production of spurious DNA (69).

### The interplay of piPolB and **Φ**29DNAP in the new piMDA protocol outperforms available MDA kits in amplifying sequences with high-GC content

As expected (28), GC bias in the amplified samples is the major limitation to homogeneous and reliable coverage and hinders the sequence assembling. Despite the low complexity of the amplified genomic sample, we found large differences in sequence coverage and assembly quality among the different WGA protocols tested. Thus, WGA amplification with the traditional random-primed MDA protocol was particularly poor for the genomes with high-GC content and promoted a very strong bias toward intermediate-GC sequences. This negative bias was only slightly reduced in the sample generated with PrimPol-MDA, with a similar pattern of overrepresented sequences. Despite a bias toward high-GC sequences, the combination of piPolB and Φ29DNAP achieved a more competent amplification of our sample, obtaining good assemblies, comparable to those of unamplified control samples.

Previous analyses reported either similar GC bias in RP-MDA and PrimPol-MDA (37) or better GC representation in PrimPol-MDA (38), using different DNA samples. Consistent with the latter, the use of T7 gp4 primase-helicase instead of random primers (pWGA) reduced GC bias and allowed the identification of high-GC species missing in MDA product, although overrepresentation of intermediate GC sequences was still similar to classical MDA (26). In this case, the authors proposed that the helicase activity of gp4 might favor the amplification of high-GC sequences (26). In the piMDA and piMDA+D methods, the priming and initial DNA synthesis are performed by the piPolB, whose strand displacement capacity might play a similar role. However, piPolB shows a strong bias towards 40-50% GC when used alone for MDA, downplaying that mechanism. Moreover, the alkaline denaturation step increases the coverage depth and breadth of all the reference genomes (Figure 6B), suggesting that high-GC content limits the initiation and also the elongation of new DNA strands by piPolB, downplaying a relevant contribution of strand displacement capacity to reduce GC bias.

Remarkably, in RP-MDA, PrimPol-MDA, and piMDA(+D) most of the amplification is performed by the same enzyme, the Φ29DNAP, suggesting that the random primers or the onset of DNA synthesis are the key factors determining the GC bias. The random primers used in MDA are usually 6-mers, and the primers of PrimPol or gp4 are also very short DNA or RNA oligonucleotides, respectively (37, 70). However, piPolB can processively synthesize longer DNA molecules of up to 6 kb (39). Conversely, greater stability of high-GC sequences would reduce processivity and possibly favor dissociation of piPolB and distributive DNA synthesis. This would generate a higher number of initiation products, albeit longer than random primers or PrimPol/gp4-synthesized primers, which in the case of piMDA(+D) could be resumed processively by the Φ29DNAP. This is consistent with the fact that the denaturation step, which would allow a higher rate of priming events by piPolB, slightly increases the GC bias in piMDA+D but not in piPolB+D (Figure 6).

Furthermore, because piPolB exhibits translesion synthesis capacity (39), it is reasonable to speculate that piPolB-mediated WGA may offer some advantages in amplifying damaged DNA, which is often present in metagenomic DNA samples. However, since piMDA uses Φ29DNAP as the main amplification enzyme, we do not expect significant differences compared with related MDA methods. Rather, an improved WGA method based only on piPolB would be more suitable for damaged DNA samples.

### Consistency of piMDA performance among duplicates and protocol modifications

The high number of reads obtained and the use of duplicates for some of the samples allowed us to analyze in detail the performance and variability of each of the amplification methods used and also the TruSeq library generation and Illumina sequencing. The overall results of the duplicate samples are very similar, although the results of the second sequencing batch resulted in a slightly lower GC bias and a better assembly that contained fewer and larger contigs and fewer misassemblies, among other parameters (Table S6-S8).

Population variant detection was also better for samples from the second sequencing batch, particularly for RP-MDA2 and piMDA+D2 samples, as compared with RP-MDA and piMDA+D, respectively. Indeed, variant calling in the piMDA+D2 sample was almost identical to the NA samples.Although conventional MDA protocols like the commercial kits tested in this work include a DNA denaturation step, we found that piPolB can perform MDA with dsDNA templates and decided to analyze the quality of the DNA products obtained with and without prior denaturation. As expected, the denaturation step increases the amount of synthesized DNA with shorter reaction times in piMDA+D compared to the piMDA protocol (Figure 3). In addition, it increases the alignment rate of reads to reference genomes (Figure 6) and improves assembly with lower N50 and L50 and fewer misassemblies (Figure 7).

However, as mentioned earlier, the denaturation step can increase GC bias (Figure 6E) and affects assembly binning (Figure 9). Therefore, unlike other available methods, piMDA can be successfully used as a simple protocol without a denaturation step or, if the amount of MDA product for downstream analysis is limiting, the prior denaturation step can be introduced. We can conclude that piMDA methods enable proficient WGA of a wide range of genomes for downstream applications, including those related to the study of microbiome diversity in different environments, especially in environments where high-GC microorganisms, such as halophiles or thermophiles, would predominate. In addition, the results shown here suggest that piMDA has great potential for application in microbiome studies involving DNA amplification, such as those using single-cell metagenomics to reconstruct strain-resolved genomes of microbial communities at once, at the risk of missing poorly represented sequences with high GC content.

## Acknowledgments

We would like to dedicate this work to the memory of Professor Margarita Salas, for her long and inspiring support and contributions to the discovery of piPolB and the early development of this project.

We also thank to all members of MRR lab for helpful discussions and critical reading of the manuscript.

