## Supplementary Figures for "A primer-independent DNA polymerase-based method for competent whole-genome amplification of intermediate to high GC sequences"

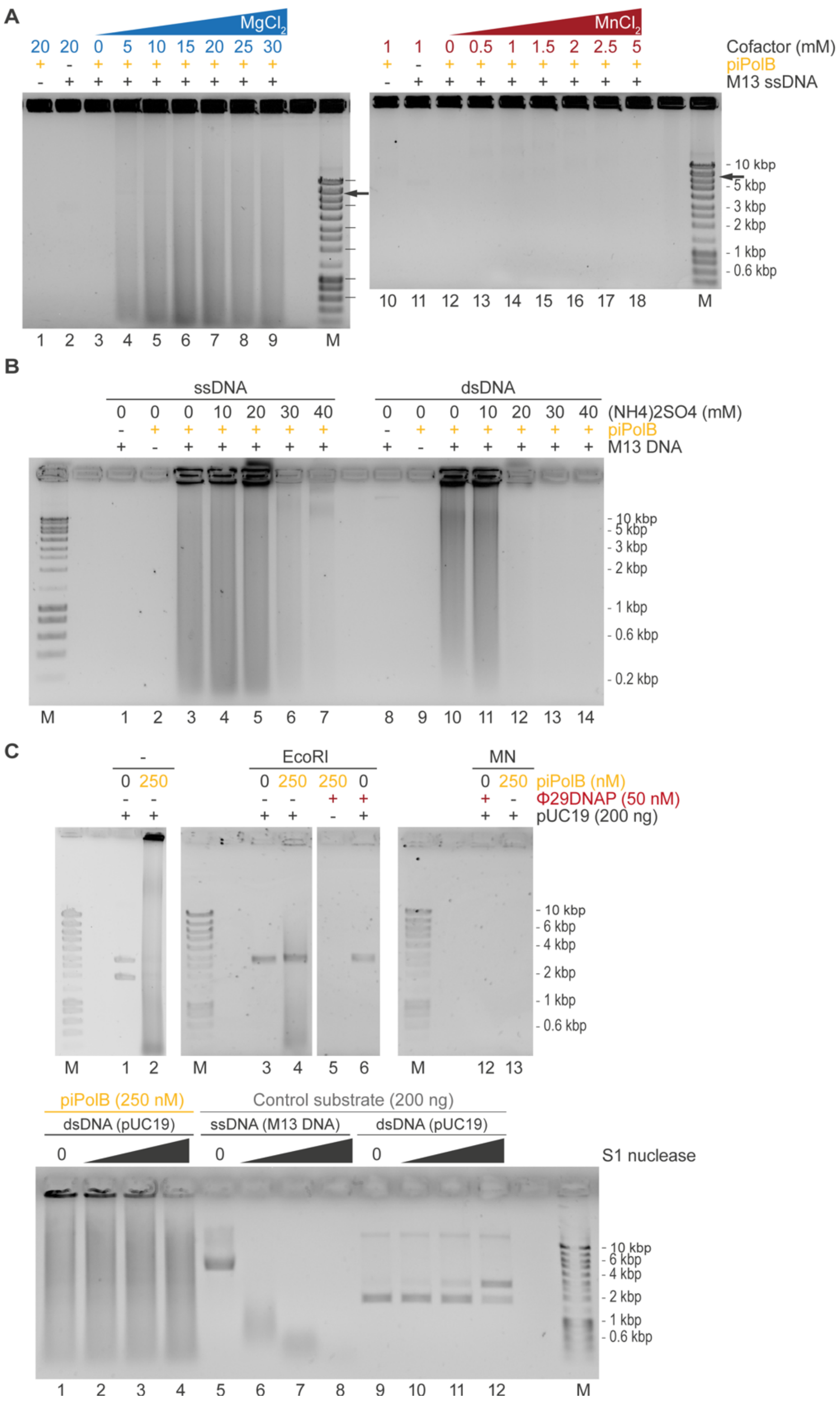


**Figure S1. Characterization of primer-independent MDA by piPolB.**

A. Agarose electrophoresis of piPolB MDA product carried out in the presence of increasing concentration of $MgCl_2$ or $MnCl_2$, as indicated.

B. Effect of ionic strength in the MDA reaction, evaluated in the presence of increasing concentration of ammonium sulfate using 20 ng of single-stranded and double-stranded M13 DNA.

C. Digestion of MDA product with specific and non-specific nucleases. Reactions were carried out with piPolB or a combination of piPolB and Φ29DNAP, as indicated. After 16 h at 30 ºC, an aliquot was withdrawn and digested with EcoRI-HF (NEB), micrococcal nuclease (MN) or single-stranded specific S1 nuclease. Control digestions on ssDNA (M13) and dsDNA plasmid (pUC19) are also shown in the bottom gel.

See Methods for details.

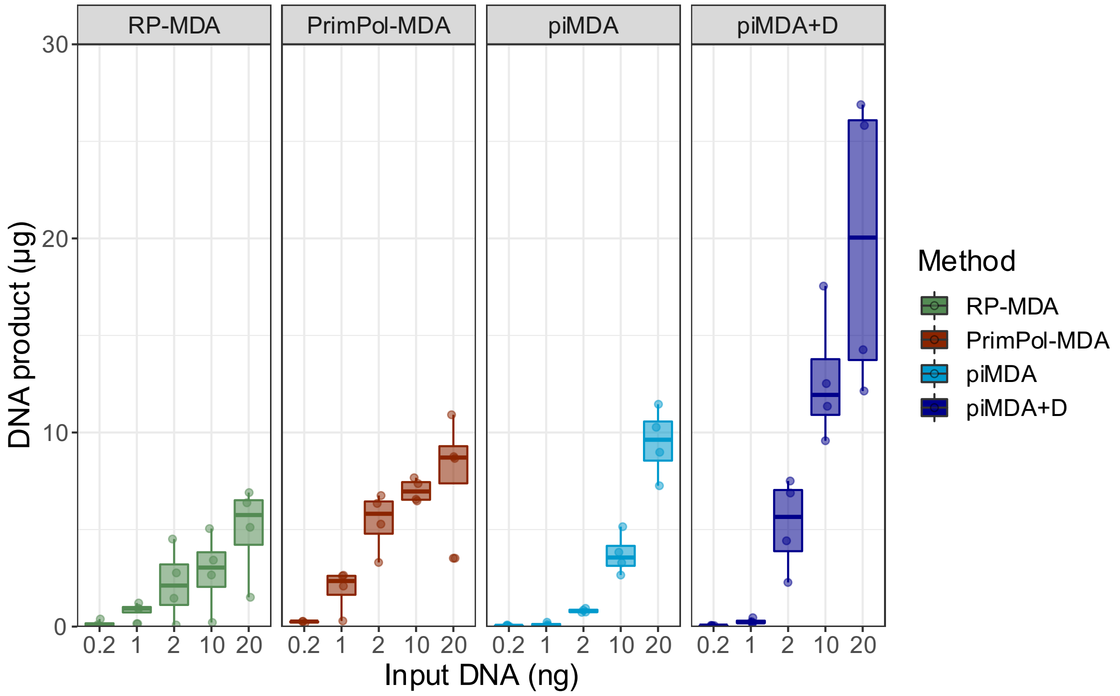


**Figure S2. Amplification sensitivity of piMDA.**
Increasing amounts of mock metagenome DNA was amplified with commercial MDA kits and piMDA or piMDA+D. Samples of four independent experiments were incubated for 3 h (PrimPol-MDA, piMDA and piMDA+D) or 16 h (RP-MDA).

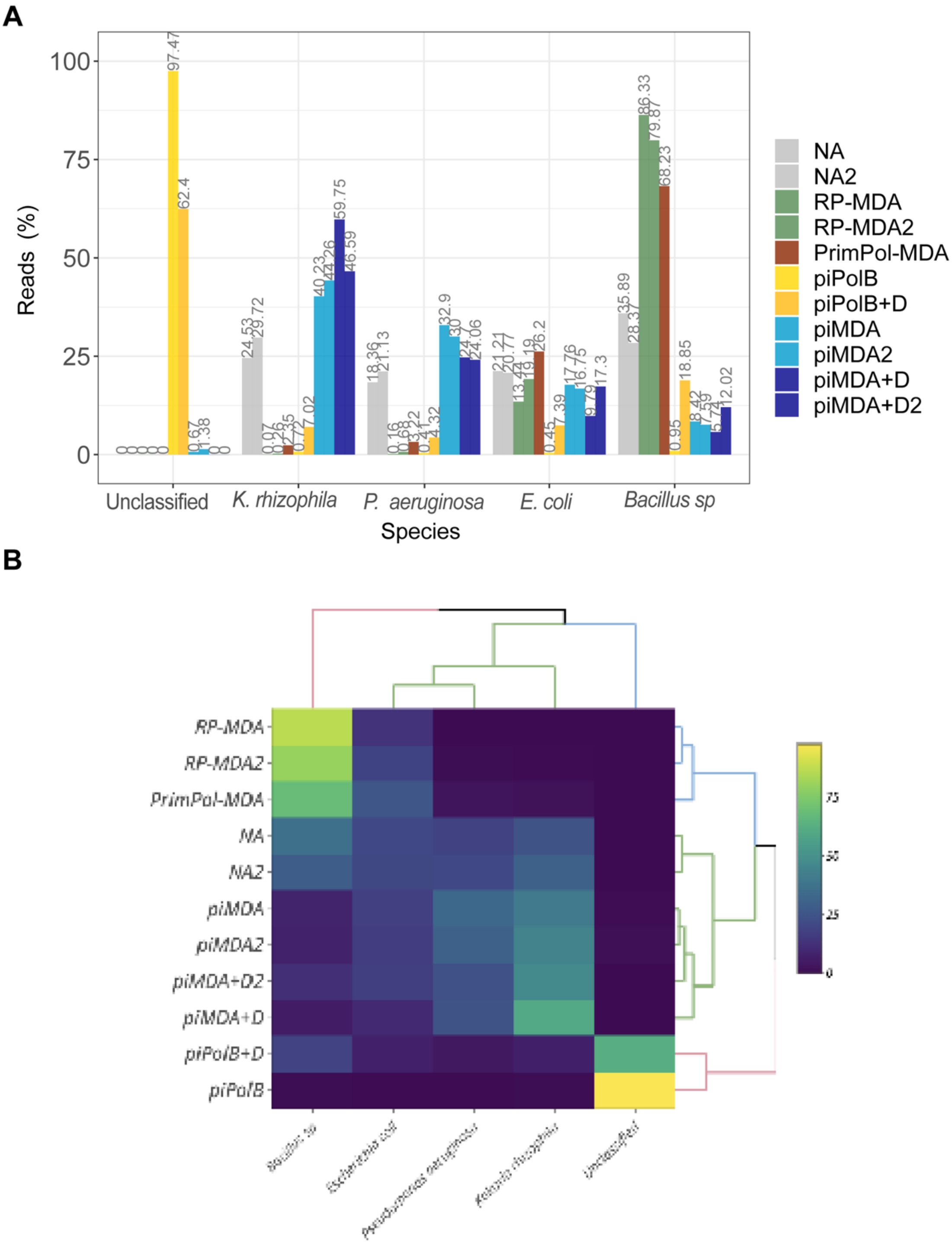


**Figure S3. Metaphlan profiling of amplified mock metagenomes sequencing.**
A. Alternative representation to Figure 4C, indicating the reads percentage assigned to each taxon.
B. Hierarchically-clustered heatmap from MetaPhlAn abundance profiles.

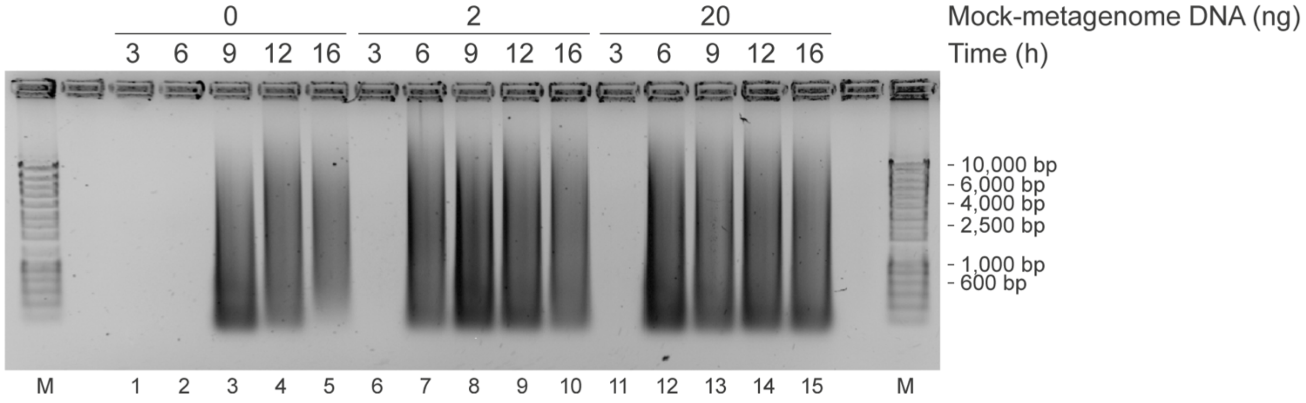


**Figure S4. *Ab initio* DNA synthesis by piPolB.**
Agarose electrophoresis of a time-course experiment of piPolB MDA in the presence or absence of input DNA substrate. Samples were incubated at 30 ºC for the indicated time.

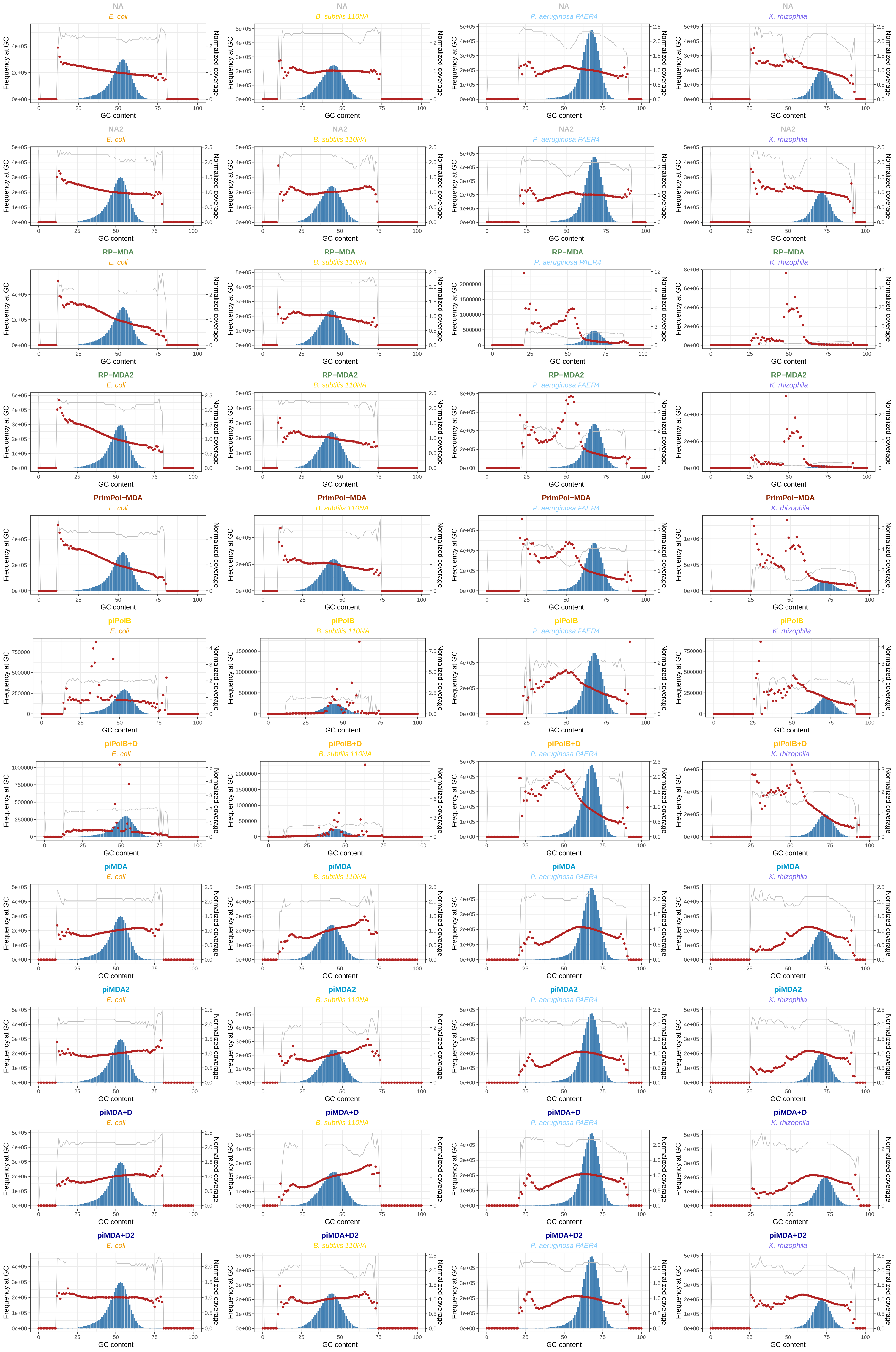

NA
E. coli
NA
B. subtilis 110NA
NA
P. aeruginosa PAER4
NA
K. rhizophila
NA2
E. coli
NA2
B. subtilis 110NA
NA2
P. aeruginosa PAER4
NA2
K. rhizophila
RP−MDA
E. coli
RP−MDA
B. subtilis 110NA
RP−MDA
P. aeruginosa PAER4
RP−MDA
K. rhizophila
RP−MDA2
E. coli
RP−MDA2
B. subtilis 110NA
RP−MDA2
P. aeruginosa PAER4
RP−MDA2
K. rhizophila
PrimPol−MDA
E. coli
PrimPol−MDA
B. subtilis 110NA
PrimPol−MDA
P. aeruginosa PAER4
PrimPol−MDA
K. rhizophila
piPolB
E. coli
piPolB
B. subtilis 110NA
piPolB
P. aeruginosa PAER4
piPolB
K. rhizophila
piPolB+D
E. coli
piPolB+D
B. subtilis 110NA
piPolB+D
P. aeruginosa PAER4
piPolB+D
K. rhizophila
piMDA
E. coli
piMDA
B. subtilis 110NA
piMDA
P. aeruginosa PAER4
piMDA
K. rhizophila
piMDA2
E. coli
piMDA2
B. subtilis 110NA
piMDA2
P. aeruginosa PAER4
piMDA2
K. rhizophila
piMDA+D
E. coli
piMDA+D
B. subtilis 110NA
piMDA+D
P. aeruginosa PAER4
piMDA+D
K. rhizophila
piMDA+D2
E. coli
piMDA+D2
B. subtilis 110NA
piMDA+D2
P. aeruginosa PAER4
piMDA+D2
K. rhizophila
Frequency at GC
Normalized coverage
GC content

**Figure S5. GC bias per reference genome.**
Normalized sequence coverage by sample reads per each reference at a given GC content was determined using Picard v2.25.0. Percentages of GC in the reference metagenome are determined from sequence bins of 100 bases, whose abundance is shown as steel blue bars (right axis). The grey line represents the average reads quality at each GC content.

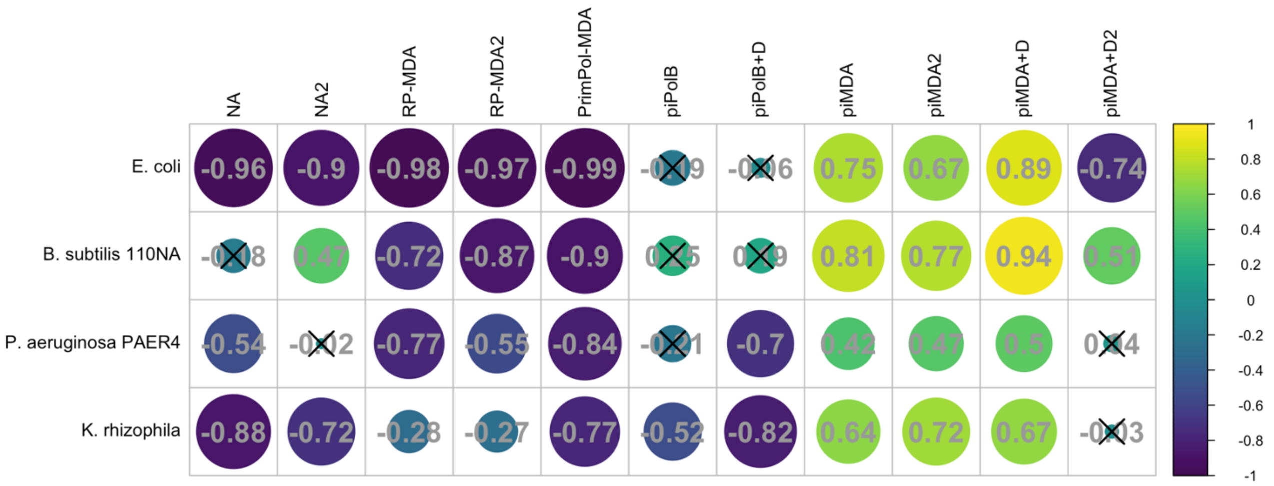


**Figure S6. Correlation of normalized coverages to sequence abundance at each GC content (95% confidence interval) per reference genome.** Nonsignificant R coefficients (p>0.05) are crossed.

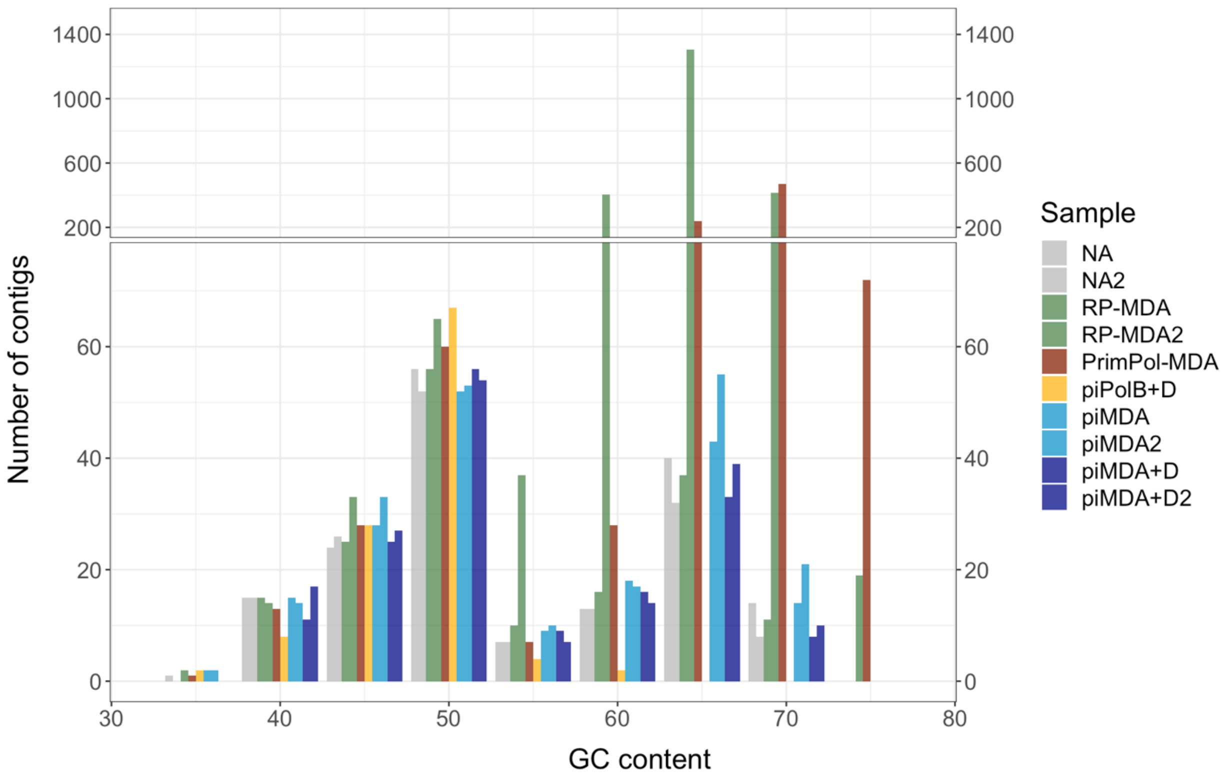


**Figure S7. Numbers of contigs per GC content.**
The number of contigs at each GC content in the reference mock metagenome was determined by Metaquast.

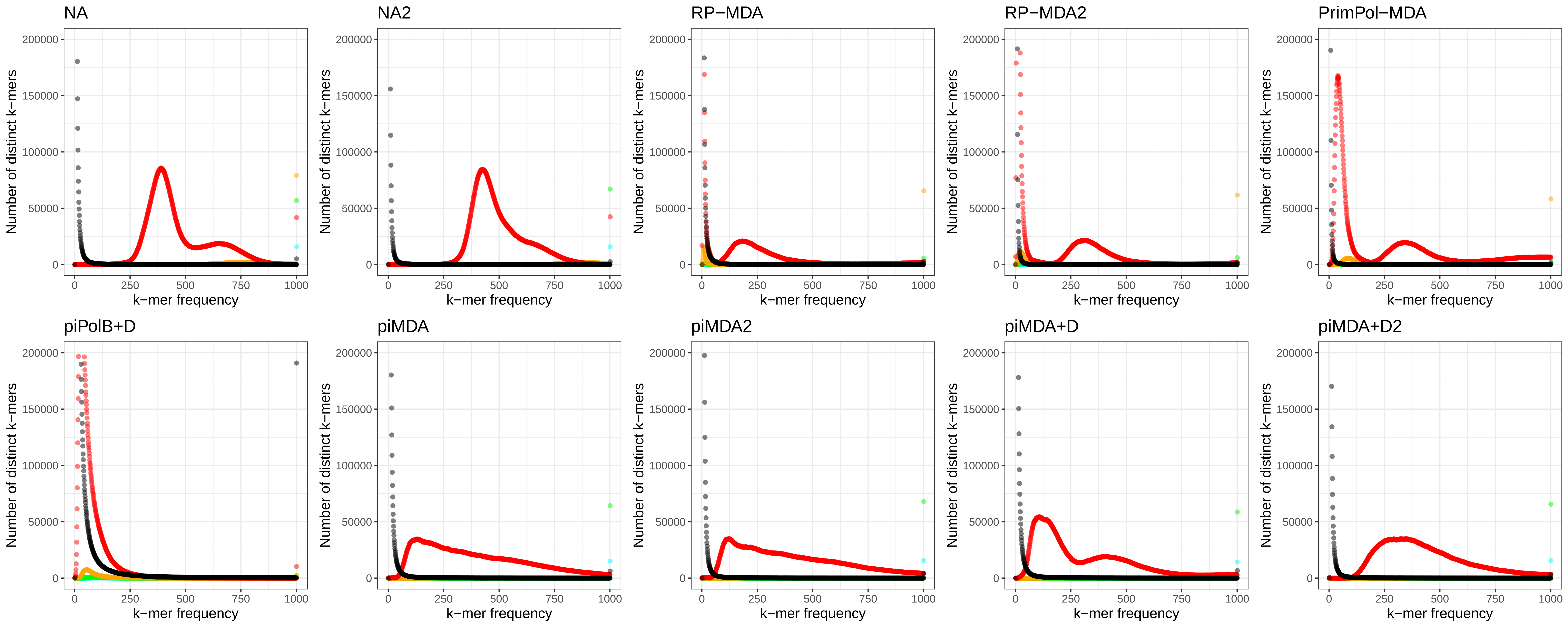

NA
NA2
RP−MDA
RP−MDA2
PrimPol−MDA
piPolB+D
piMDA
piMDA2
piMDA+D
piMDA+D2
Number of distinct k−mers
k−mer frequency
0
50000
100000
150000
200000
0
250
500
750
1000

**Figure S8. Assembly spectra copy number plots.**
KAT comparative spectra were used to check assemblies' coherence against the content within reads that were used to produce each assembly. Color lines represent k-mer content presence 0 (black), 1 (red), 2 (orange) or <3 (cyan) times in the assembly.
